# An unexpected role for the conserved ADAM-family metalloprotease ADM-2 in *Caenorhabditis elegans* molting

**DOI:** 10.1101/2021.12.15.472830

**Authors:** Braveen B. Joseph, Sarina Meadows, Phillip T. Edeen, Shaonil Binti, David S. Fay

## Abstract

Molting is a widespread developmental process in which the external extracellular matrix (ECM), the cuticle, is remodeled to allow for organismal growth and environmental adaptation. Studies in the nematode *Caenorhabditis elegans* have identified a diverse set of molting-associated factors including signaling molecules, intracellular trafficking regulators, ECM components, and ECM-modifying enzymes such as matrix metalloproteases. *C. elegans* NEKL-2 and NEKL-3, two conserved members of the NEK family of protein kinases, are essential for molting and promote the endocytosis of environmental steroid-hormone precursors by the epidermis. Steroids in turn drive the cyclic induction of many genes required for molting. Here we report a novel role for the sole *C. elegans* ADAM–meltrin metalloprotease family member, ADM-2, as a negative regulator of molting. Whereas loss of *adm-2* led to strong suppression of molting defects in partial loss-of-function *nekl* mutants, overexpression of ADM-2 induced molting defects in wild-type animals. CRISPR genome editing implicated the Zn-binding motif within the metalloprotease domain as critical for mediating the effects of ADM-2 on molting. ADM-2 is expressed in the epidermis, and its trafficking through the endo-lysosomal network was disrupted after NEKL depletion. We also identified the epidermally expressed low-density lipoprotein receptor–related protein, LRP-1, as a candidate target of ADM-2 regulation. Whereas loss of ADM-2 activity led to the upregulation of LRP-1, ADM-2 overexpression caused a reduction in LRP-1 abundance, suggesting that ADM-2 may function as a sheddase for LRP-1. We propose that loss of *adm-2* suppresses molting defects in *nekl* mutants by eliminating a negative regulator of LRP-1, thereby compensating for defects in the efficiency of LRP-1 and sterol uptake. Our findings emphasize the importance of endocytic trafficking for both the internalization of factors that promote molting and the removal of proteins that would otherwise be deleterious to the molting process.

**Author Summary:** The molecular and cellular features of molting in nematodes share many similarities with cellular and developmental processes that occur in mammals. This includes the degradation and reorganization of extracellular matrix materials that surround cells, as well as the intracellular machineries that allow cells to communicate and sample their environments. In the current study, we found an unexpected function for a conserved protein that degrades proteins on the external surface of cells. Rather than promoting molting through extracellular matrix reorganization, the ADM-2 protease can inhibit the molting process. This observation can be explained by data showing that ADM-2 negatively regulates LRP-1, a membrane protein that promotes molting by facilitating the uptake of molecular building blocks at the cell surface that are needed for molting-related signaling. Our data provide insights into the mechanisms controlling molting and link several conserved proteins to show how they function together during development.

## Introduction

The cuticle of *Caenorhabditis elegans* is an external extracellular matrix (ECM) required for locomotion, body shape maintenance, and protection from the environment [1, 2]. During larval development *C. elegans* undergoes four molts, a specialized form of apical ECM remodeling, whereby a new cuticle is synthesized under the old cuticle, which is partially degraded and shed [1, 2]. *C. elegans* molting cycles are orchestrated by conserved steroid-hormone receptors, including NHR-23 (an ortholog of human RORC) and NHR-25 (an ortholog of human NR5A1), which collectively control the oscillation of hundreds of mRNAs [2–4]. The production of molting-specific steroid-hormone ligands is thought to be dependent in part on the uptake of environmental sterols by epidermally expressed LRP-1 (the homolog of human LRP2/megalin), a member of the low-density lipoprotein receptor–related protein family [5, 6]. Consistent with this, internalization of LRP-1 by clathrin-mediated endocytosis (CME) is essential for normal molting [2, 5–7].

We have previously shown that the *C. elegans* protein kinases NEKL-2 (an ortholog of human NEK8/9) and NEKL-3 (an ortholog of human NEK6/7) promote endocytosis of LRP-1 at the apical epidermal plasma membrane. Correspondingly, loss of either NEKL-2 or NEKL-3 function leads to a reduction or delay in the expression of molting genes, a failure to complete molting, and larval arrest [2, 8–12]. NEKL-2 and NEKL-3 (NEKLs) are members of the NIMA-related kinase (NEK) protein family, mammalian orthologs of which have been studied primarily in the context of cell cycle regulation and ciliogenesis [13–28]. *C. elegans* NEKLs bind to and co-localize with several conserved ankyrin-repeat proteins, MLT-2 (an ortholog of human ANKS6), MLT-3 (an ortholog of human ANKS3), and MLT-4 (an ortholog of human INVS), referred to here collectively as MLTs, which are essential for the proper localization of NEKLs [9]. Correspondingly, loss of MLT functions leads to molting defects that are identical to those observed with loss of the NEKLs [9]. NEKLs and MLTs form two distinct complexes (NEKL-2– MLT-2–MLT-4 and NEKL-3–MLT-3) and are expressed in a punctate pattern in the large epidermal syncytium known as hyp7, in which they are specifically required [2, 8, 9].

The cellular and physiological mechanisms by which NEKLs–MLTs impact the molting process through intracellular trafficking have yet to be fully explored. It is likely that NEKLs are required for the uptake and processing of membrane cargo, including LRP-1, that is required for molting. Using a forward-genetics suppressor approach [29], we previously found that loss of AP2 clathrin-adapter subunits, as well as the AP2 allosteric activator FCHO-1 can suppress molting and trafficking defects in NEKL mutants [11]. These and other studies revealed that NEKLs control endocytosis in part by facilitating the uncoating of sub-apical clathrin-coated vesicles and may affect additional trafficking processes through the regulation of actin via the CDC-42–SID-3 (corresponding to the human CDC42–ACK1/2) pathway [10, 11].

Here we report suppression of *nekl* molting defects caused by loss of the conserved ADM-2 transmembrane metalloprotease. Curiously, although proteases have previously been implicated as positive regulators of molting, we find that ADM-2 exerts a negative influence on the molting process. ADM-2 belongs to the ADAM (<u>a</u> <u>d</u>isintegrin <u>a</u>nd <u>m</u>etalloprotease) family of metallopeptidases, which are members of the zinc protease superfamily [30, 31]. ADM-2 is the sole member of the meltrin metalloprotease subfamily in *C. elegans* [32, 33], which in humans consists of ADAM9 (Meltrin γ), ADAM12 (Meltrin α), ADAM19 (Meltrin β), and ADAM33 [34, 35]. Notably, knockouts of meltrin family members in mammals have generally not provided clear insights into the roles of meltrins in vivo during development, which may in part be due to genetic redundancies [32, 36, 37]. Here we show that, unlike AP2 and *fcho-1* mutants, loss of ADM-2 function did not suppress *nekl-*associated trafficking defects. Rather, ADM-2 was itself dependent on NEKL–MLTs for its uptake and endocytic processing from the epidermal surface where it may act as a negative regulator of LRP-1. Our findings further suggest that loss of ADM-2 may specifically bypass trafficking defects in weak loss-of-function *nekl* mutants by de-repressing LRP-1. Thus, NEKLs may be required for both the internalization of positive and negative regulators of the molting cycle.

## Results

### *nekl* molting defects are suppressed by reduced function of the ADM-2 metalloprotease

We previously described an approach to identify genetic suppressors of partial loss-of-function mutations in NEKL kinases [29]. Whereas strains homozygous for either *nekl-2(fd81)* or *nekl-3(gk894345)* weak loss-of-function alleles are viable, *nekl-2(fd81)*; *nekl-3(gk894345)* double mutants (hereafter referred to as *nekl-2; nekl-3* mutants) are synthetically lethal and exhibit ~98.5% larval arrest due to L2/L3 molting defects [9]. In the absence of a suppressor mutation, propagation of *nekl-2; nekl-3* mutants requires the presence of a GFP-marked *nekl-2+* or *nekl-3+* transgenic rescuing array (Fig 1A and 1E). In contrast, strains homozygous for the suppressor alleles *fd130* or *fd163* of are ~80% viable and propagate in the absence of a rescuing array (Fig 1B, 1C, and 1E).

**Fig 1.**
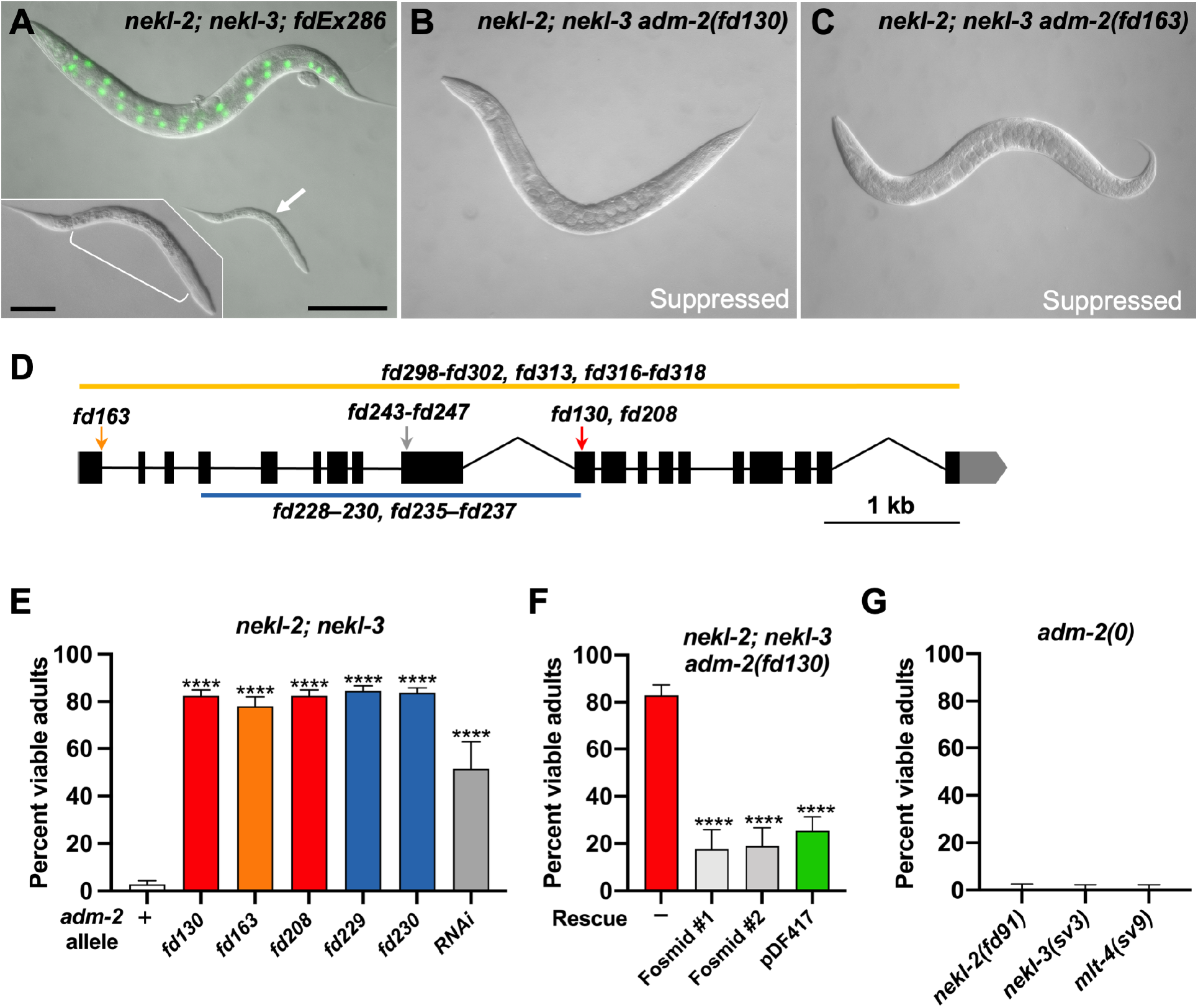
Loss of ADM-2 function suppresses *nekl* molting defects. (A) DIC image of *nekl-2; nekl-3* double-mutant worms. The adult worm contains a rescuing extrachromosomal array *(nekl-3*^*+*^*; sur-5::GFP)*. An arrested larva is marked by the white arrow and enlarged in the inset; the white bracket indicates the constricted region containing a double cuticle. (B, C) DIC images of *nekl-2; nekl-3* double-mutant adult worms containing the *adm-2(fd130)* (B) and *adm-2(fd163)* (C) mutant alleles. Bar size in A = 100 μm (for A–C); in inset, 20 μm. (D) Schematic diagram of the *adm-2* locus. Solid black rectangles indicate exons; introns are demarcated by black lines. Locations of the *fd163*, *fd130*, *fd208*, and *fd243–fd247* alleles are indicated by arrows. Large deletion alleles *fd298–fd302 fd313, fd316–fd318, fd228–fd230*, and *fd235–fd237* are indicated by orange and blue lines. (E) Bar plot showing percentage of viable adult-stage *nekl-2; nekl-3* worms with the indicated *adm-2* alleles (or RNAi); + indicates wild-type *adm-2*. (F) Bar plot showing reversion of suppression in *nekl-2; nekl-3 adm-2(fd130)* mutants by fosmids expressing wild-type *adm-2* and by an *adm-2* cDNA fused to GFP (pDF417). Fosmid #1 and #2 indicate two independent extrachromosomal lines. (G) Bar plot showing failure to suppress molting defects in *nekl–mlt* hypomorphic mutants by *adm-2* null mutants [*nekl-2(fd91); adm-2(fd313)*, *nekl-3(sv3); adm-2(fd316)*, and *mlt-4(sv9); adm-2(fd317)*]. Error bars in E–G represent 95% confidence intervals. p-Values were determined using Fisher’s exact test; ****p ≤ 0.0001.

Using our protocols for whole-genome sequencing and bioinformatical analysis [29], we identified the causal mutation corresponding to *fd130* to be a G-to-A transition in exon 10 of *adm-2*/C04A11.4 (Fig 1D). *fd130* converts codon 494 (TG<u>G</u>; W) into a premature translational termination signal (TG<u>A</u>; stop codon), resulting in the predicted truncation of the 952-amino-acid protein. Correspondingly, the independently isolated allele *fd163* is a G-to-A transition in the conserved 5’ splice donor sequence in the first intron of *adm-2* (<u>G</u>T to <u>A</u>T) (Fig 1D). The *fd163* mutation is predicted to result in a stop codon immediately following R66. Using CRISPR/Cas9 methods we isolated several additional *adm-2* alleles including *fd208*, a 1-bp deletion that causes a frameshift after Y479, along with *fd229* and *fd230*, deletions that span exons 4–10 and that result in frame shifts after S123 and T122, respectively (Fig 1D). Like *fd130* and *fd163*, these alleles led to similarly robust suppression of molting defects in the *nekl-2; nekl-3* background and are predicted to result in strong or complete loss of ADM-2 function (Fig 1E).

Several additional pieces of evidence indicate that it is loss of ADM-2 function that leads to suppression of *nekl-2; nekl-3* molting defects. (1) *adm-2* mutations that suppress these molting defects (e.g., *fd130* and *fd163*) are fully recessive (see Materials and Methods). (2) Extrachromosomal expression of a fosmid clone containing wild-type genomic ADM-2 sequences strongly reversed suppression in *nekl-2; nekl-3 adm-2* mutants (Fig 1F). (3) RNAi of *adm-2* led to significant suppression of molting defects in *nekl-2; nekl-3* mutants (Fig 1E). We note that loss of *adm-2* in wild-type backgrounds, including a strong loss-of-function deletion allele, *fd300*, did not appear to impair development, health, or fecundity, indicating that *adm-2* is a non-essential gene (S1A Fig). Consistent with this, no phenotypes have been previously ascribed to *adm-2* mutations.

ADM-2 is a member of the ADAM (<u>a</u> <u>d</u>isintegrin <u>a</u>nd <u>m</u>etalloprotease) family of metallopeptidases, with its closest human homologs belonging to the meltrin subfamily (ADAM9/12/19/33) (S1C Fig) [32, 34, 35]. Meltrins are notable for having functional proteases that contain a histidine-coordinated zinc-binding site, which is also found in ADM-2 (S2 Fig). Like other meltrins, ADM-2 contains an N-terminal cysteine switch, cysteine loop, and disintegrin domain; a transmembrane domain; and several predicted SH3-binding sites in its cytoplasmic C terminus (S2 Fig) [31, 38]. Although linked to a range of human diseases, individual loss-of-function mutations in mouse meltrins have generally not produced robust developmental defects, and no phenotypes have been associated with either of the two *Drosophila* meltrin family members [32, 36, 37].

We note that WormBase annotates two *adm-2* isoforms that are identical through exon 18 (corresponding to aa A915) but differ at the 5’ ends of their 19^th^ (terminal) exons; *adm-2a* and *adm-2b* are predicted to encode 952 and 929 aa proteins, respectively. The noncanonical 18th intron acceptor site of *adm-2b* (5-‘GCAAAAG-3’) occurs 7 bp upstream of the corresponding acceptor site of *adm-2a* (5’-ATTTCAG-3’) and terminates translation 76 bp upstream of the stop codon of *adm-2a*. The peptide regions corresponding to exon 19 of *adm-2a* (37 aa) and *adm-2b* (14 aa) do not contain any known domains nor were homologies detected with other proteins.

To determine if other *C. elegans* ADAM family members may also contribute to molting control, we tested ten other family members for their ability to suppress *nekl-2; nekl-3* molting defects (S1B Fig). We failed to detect suppression after inhibition of each gene using RNAi (dsRNA) injection methods, which were effective in promoting suppression when targeting *adm-2*. Thus, the suppression of *nekl*-associated molting defects by *adm-2* is unique among the *C. elegans* ADAM family members.

### Suppression by ADM-2 occurs via a mechanism that is distinct from trafficking suppressors

Loss of AP2 clathrin-adapter subunits and loss of the AP2 activator FCHO-1 individually suppress strong and/or null mutations in NEKLs and MLTs through their effects on CME [11]. We therefore tested if an *adm-2* null mutation could also suppress strong loss-of-function alleles of *nekls* and *mlts*. Notably, loss of *adm-2* was unable to restore viability to *nekl-2(fd91)*, *nekl-3(sv3)*, or *mlt-4(sv9)* strong loss-of-function alleles, which typically arrest as partially constricted L2/L3 larvae (Fig 1G). These results suggest that the mechanisms underlying the suppression of molting defects by *adm-2* and CME-associated trafficking factors may be distinct.

To directly test if loss of ADM-2 can suppress CME defects in *nekls*, we examined the localization of GFP-tagged LRP-1/megalin, an apical transmembrane cargo that is trafficked via CME. Using the auxin-inducible degron (AID) system to remove NEKL-3 activity in 1-day-old adult worms [11], we observed a dramatic accumulation of LRP-1::GFP at or near the apical membrane (Fig 2A and 2D), consistent with our previous findings [11]. As anticipated, loss of FCHO-1 partially corrected LRP-1::GFP mislocalization defects in NEKL-3::AID-depleted worms (Fig 2B and 2D), consistent with the ability of *fcho-1* mutations to suppress *nekl*-associated clathrin localization and mobility defects [11]. In contrast, complete loss of ADM-2 failed to correct LRP-1::GFP defects in NEKL-3::AID-depleted adults and if anything showed slightly enhanced apical LRP-1::GFP accumulation relative to the NEKL-3::AID-depleted worms (Fig 2C and 2D; also see below). Collectively, these results indicate that ADM-2 does not suppress *nekl* molting defects by correcting CME deficits and is therefore likely to act through a distinct mechanism.

**Fig 2.**
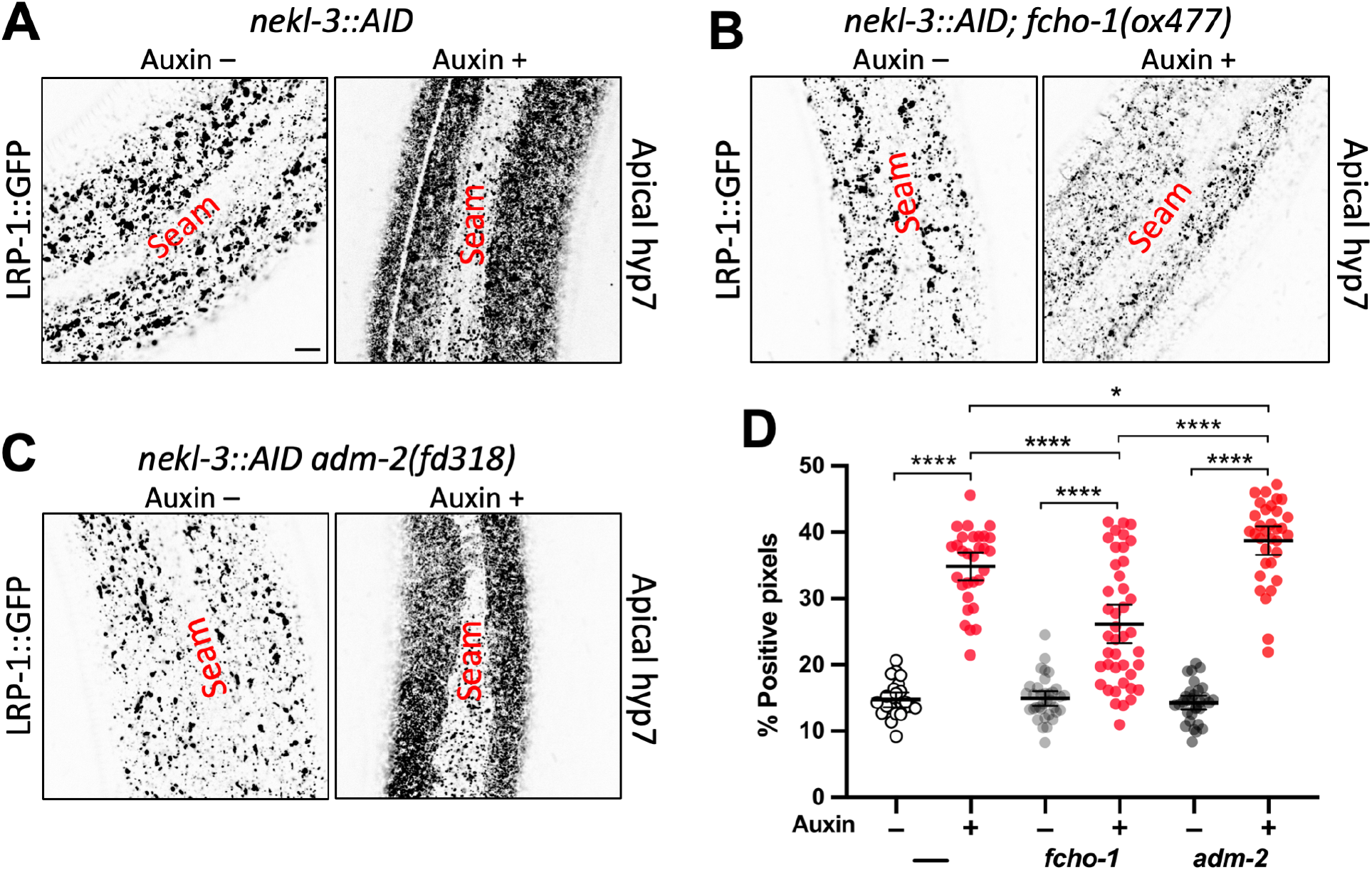
Loss of *adm-2* function does not correct the *nekl* trafficking defects. (A–C) Representative confocal images of 2-day-old adult worms expressing LRP-1::GFP in the apical hyp7 region of the epidermis. LRP-1 expression in the *nekl-3::AID* (A), *nekl-3::AID*; *fcho-1(ox477)* (B), and *nekl-3::AID; adm-2(fd318)* (C) mutant backgrounds in the absence (–) and presence (+) of auxin treatment. Bar in A = 5 μm (for A–C). (D) Dot plot showing the percentage of GFP-positive pixels within the apical plane of the worm epidermis for individuals of the specified genotypes and auxin treatment groups. Group means along with 95% confidence intervals (error bars) are indicated. p-Values were obtained by comparing means using an unpaired t-test: ****p ≤ 0.0001, *p ≤ 0.05.

### ADM-2 is expressed in multiple tissues including the epidermis

To gain insight into how ADM-2 may affect molting in *nekl* mutants, we examined endogenously tagged *adm-2::mScarlet* and *adm-2::GFP* strains, in which the fluorescent marker was fused to the C terminus of the ADM-2a cytoplasmic domain. Both CRISPR-tagged versions showed a punctate pattern within hyp7, a large epidermal syncytium that encompasses most of the central body region of the worm, including localization to small puncta near the apical membrane (Fig 3A–E’). Notably, both NEKLs and MLTs are specifically expressed and required in the hyp7 syncytium [8, 9]. We also detected some differences between the localization patterns of ADM-2::mScarlet and ADM-2::GFP. In particular, ADM-2::mScarlet was observed in larger vesicular and tubular-like structures throughout the epidermis, whereas these structures were mostly absent in ADM-2::GFP worms (Fig 3A–D; also see below).

**Fig 3.**
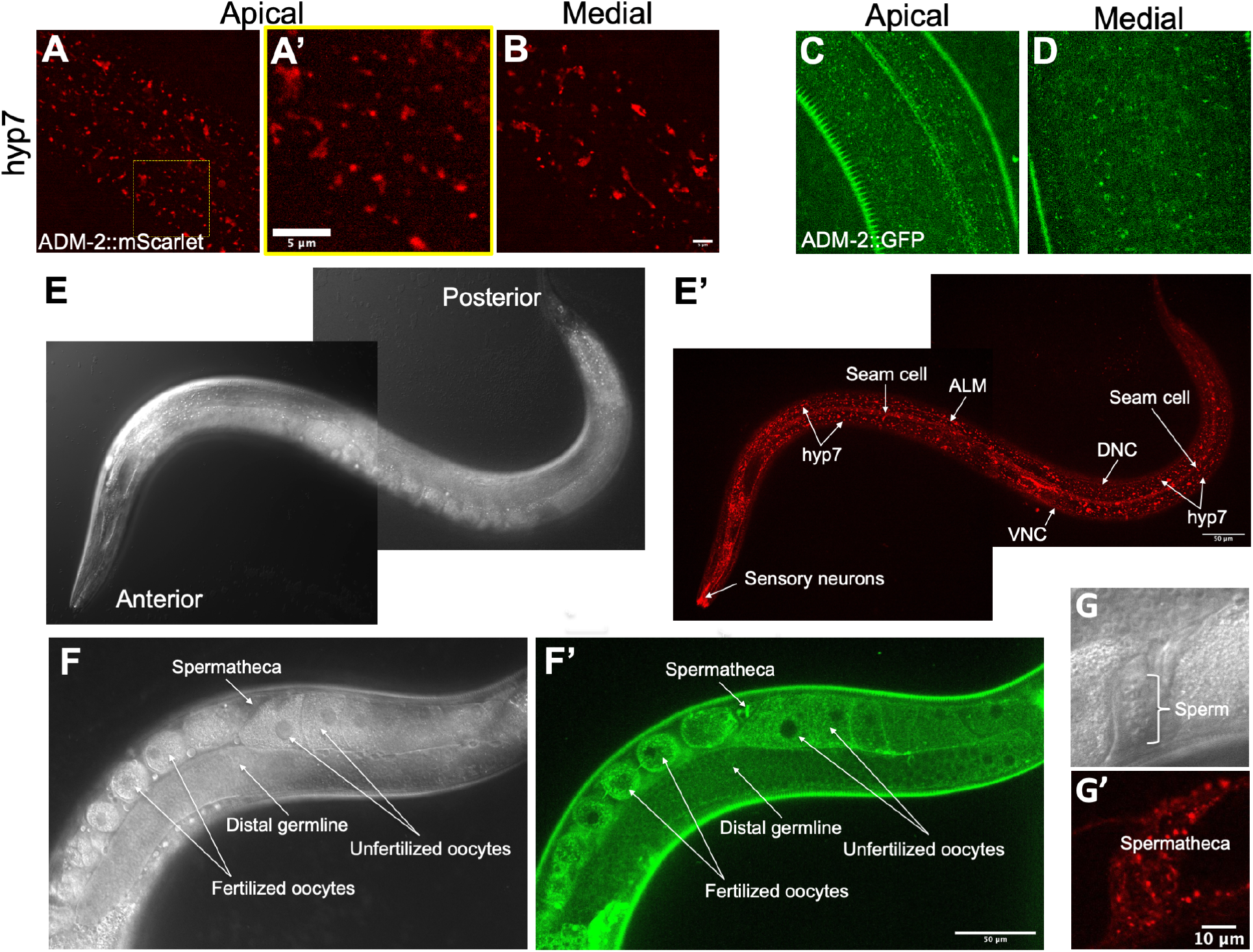
Expression of ADM-2 in the epidermis and other tissues. (A–D and A’) Representative confocal images of ADM-2 expression in the *C. elegans* hyp7 region. ADM-2::mScarlet (A, B) and ADM-2::GFP (C, D) expression at apical (A, C) and medial (B, D) planes. A’ is the inset from panel A. Bar in B = 5 μm (for A–D); in inset A’ = 5 μm. (E–G and E’–G’) Representative DIC (E–G) and confocal (E’–G’) images of ADM-2 expression. ADM-2::mScarlet in the hyp7 hypodermis; seam cells; sensory neurons; and ALM, DNC, and VNC neurons (E, E’) and in spermatheca (G, G’). ADM-2::GFP expression in the distal germline, oocytes, and spermatheca (F, F’). Bar in E’ = 50 μm (for E, E’); in G’ = 10 μm (for G, G’); in F’ = 50 μm (for F, F’).

ADM-2 was also detectable in seam cells, a lateral epidermal syncytium that borders hyp7 along the length of the animal (Fig 3E and 3E’); in the anterior epidermis (S3A and A’); and in a variety of head, tail, and centrally located neurons (Fig 3E and 3E’; S3B–C and S3B’–C’ Fig). In addition, ADM-2 was observed in proximal oogenic cells of the hermaphrodite germline, with levels increasing in maturing oocytes, where it was localized to the cytoplasm and plasma membrane (Fig 3F and 3F’). Likewise, ADM-2 is expressed in fertilized oocytes (Fig 3F and 3F’) and throughout embryogenesis (S3D–F and S3D’–F’ Fig). In contrast, ADM-2 was not detected in mature sperm cells but was expressed in myoepithelial cells of the hermaphrodite spermatheca (Fig 3G and 3G’). We also note that ADM-2::GFP expression using a multicopy reporter under the control of the *adm-2* promoter region showed strongest expression in neurons where it accumulated at or near the plasma membrane (S3D Fig). These findings are consistent with ADM-2 acting in the epidermis to affect molting, but they also suggest that ADM-2 may have functions in other tissues.

### ADM-2 is trafficked through endo-lysosomal compartments and is sensitive to NEKL activities

To determine the identity of ADM-2 puncta, vesicles, and tubular-like structures in the epidermis, we performed colocalization experiments first using a CRISPR-tagged clathrin heavy chain marker, GFP::CHC-1 [11]. Although statistically insignificant, rare colocalization between ADM-2::mScarlet and apical clathrin was occasionally detected (Fig 4A–C’, S4A Fig). We note that endogenous plasma membrane–localized ADM-2::GFP and ADM-2::mScarlet both presented with extremely faint signals within hyp7 (Fig 4A and 4C), making detection and colocalization of this population difficult to assay; this suggests that ADM-2 may be rapidly turned over at the plasma membrane, either by CME or through a CME-independent mechanism. We note that mammalian ADAMs, including the meltrin family, are internalized via CME [39–42]. In addition, we observed little or no colocalization between ADM-2::mScarlet and medial GFP::CHC-1 clathrin-containing structures (S4C–E and S4C’–E’ Fig), which may represent clathrin-coated vesicles emanating from the trans Golgi or recycling endosomes.

**Fig 4.**
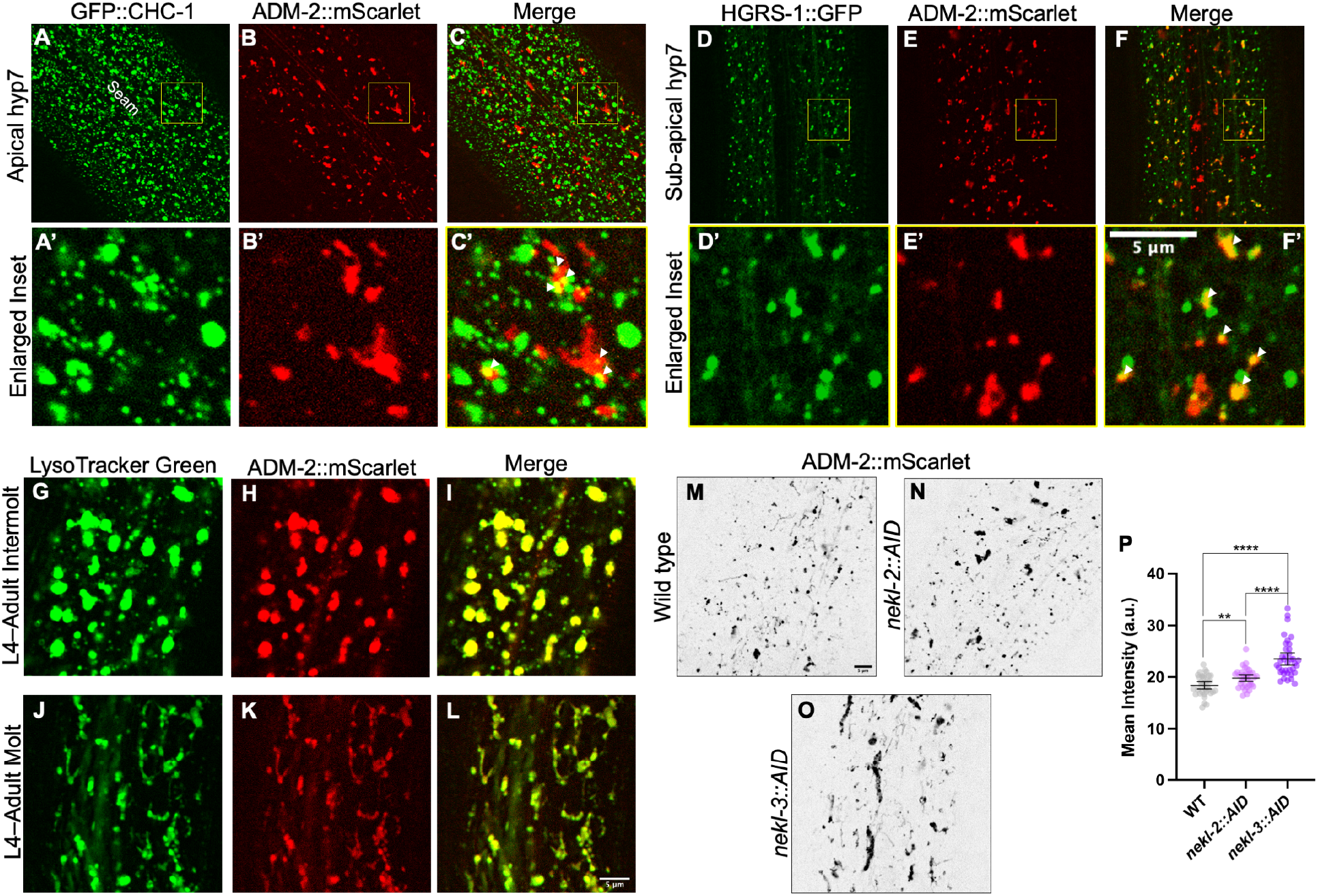
ADM-2 localization to endocytic compartments is affected by NEKL activity. (A–L) Co-localization analysis of ADM-2::mScarlet with GFP::CHC-1 (A–C), Phyp7::HGRS-1::GFP (D–F), and LysoTracker Green (G– L) within apical (A–C) and sub-apical or medial (D–L) planes of hyp7. A’–F’ are insets of A–F confocal images. White arrowheads in C’ and F’ indicate colocalized puncta. For LysoTracker studies (G–L), representative confocal images during intermolt (G–I) and molting (J–L) stages are shown. (M–P) Apical hyp7 ADM-2::mScarlet localization in auxin-treated wild-type (M), *nekl-2::AID* (N), and *nekl-3::AID* (O) 2-day-old adults with average mean intensity calculations (P). Group means along with 95% confidence intervals (error bars) are indicated. p-Values were obtained by comparing means using an unpaired t-test: ****p ≤ 0.0001, **p ≤ 0.01. Bar in M = 5 μm (for A–F and M–O); in F’ = 5 μm (for insets A’–F’); in L = 5 μm (for G–L).

In contrast, ADM-2::mScarlet exhibited partial colocalization with the endosomal marker P_hyp7_*::hgrs-1::GFP* in both the sub-apical and medial planes (Fig 4D–F’, S4A–B, S4F–H and S4F’– H’ Fig). HGRS-1/HRS localizes to early endosomes and multivesicular bodies and is a component of the ESCRT-0 complex, which, together with ESCRT-I–III, promotes cargo sorting and lysosomal targeting [43–46]. Consistent with this, medial ADM-2::mScarlet showed strong co-localization within the lysosomal marker LysoTracker Green during intermolts (Fig 4G–I), when lysosomes appear roughly spherical, and during molting periods (Fig 4J–L), when lysosomes acquire a tubular morphology [47]. Thus, following rapid uptake into endosomes, ADM-2 is likely degraded by lysosomes, although some portion may be recycled back to the plasma membrane. Degradation of ADM-2 by lysosomes is further supported by the relative absence of medial ADM-2::GFP accumulation (Fig 3D), as GFP is acid sensitive and fluorescence is rapidly quenched within maturing endosomes and lysosomes [47, 48].

Given our previous observations showing that NEKL–MLT proteins are required for normal trafficking within hyp7 [8, 10, 11], we tested if depletion of NEKLs caused changes in the abundance and subcellular localization of ADM-2. Notably, we observed total levels of ADM-2::mScarlet to increase slightly in auxin-treated *nekl-2::AID* adults, with more robust changes occurring in auxin-treated *nekl-3::AID* animals (Fig 4M–P). Consistent with this, partial knockdown of MLT-3 in adults by RNAi led to modest increases in the levels of ADM-2::mScarlet and GFP::CHC-1 (S5A–B Fig). In worms that had undergone NEKL::AID depletion, ADM-2::mScarlet increased slightly in total levels and accumulated in large internal endocytic or lysosome-like structures (Fig 5M–P). Collectively, these findings indicate that NEKL activities impact ADM-2 trafficking within hyp7.

**Fig 5.**
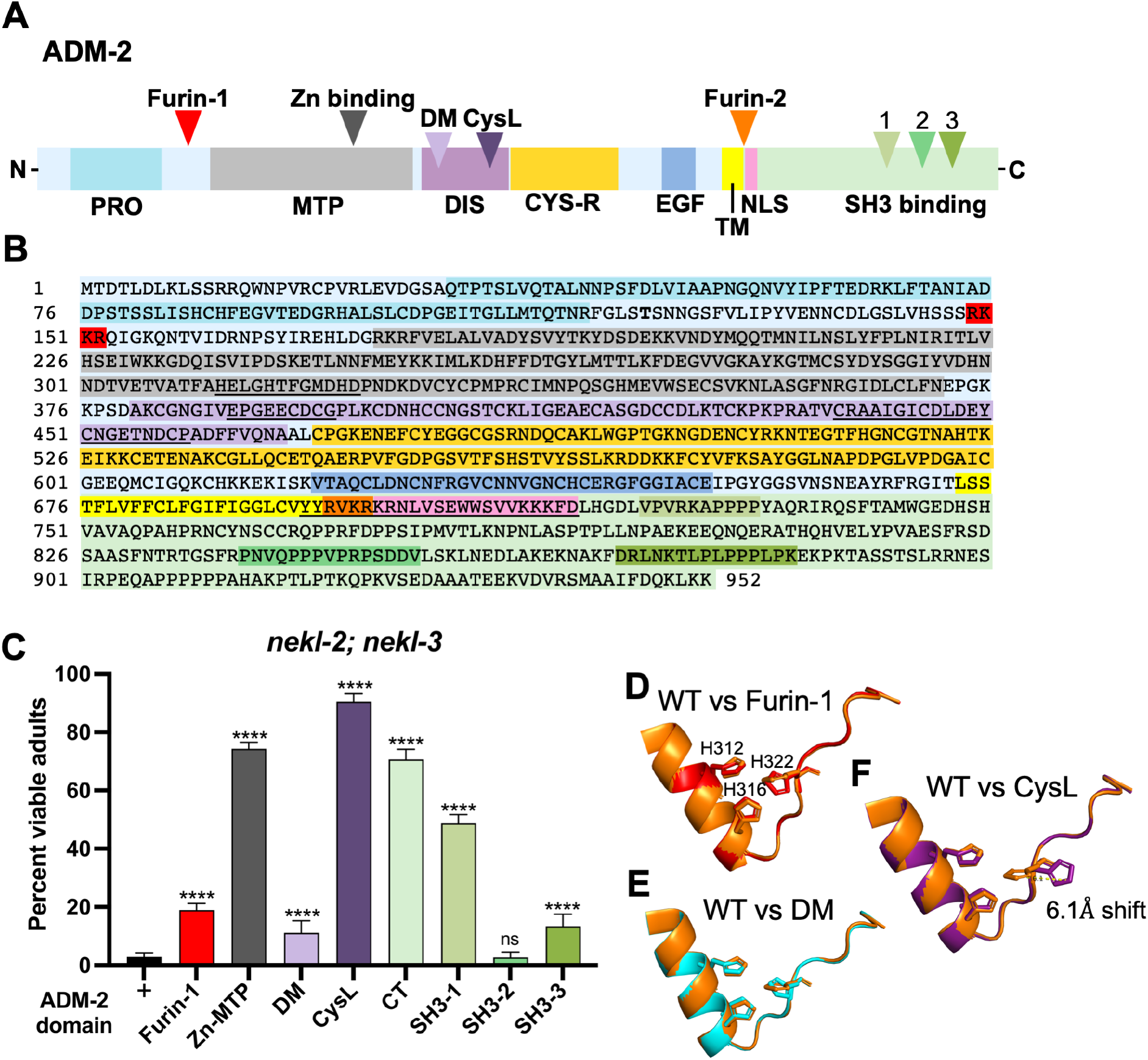
Functional analysis of ADM-2 domains. Schematic representation of predicted protein domains within ADM-2. PRO, prodomain; MTP, metalloprotease domain; DIS, disintegrin domain; CYS-R, cysteine repeat region; TM, transmembrane domain; NLS, predicted nuclear localization signal; DM, disintegrin motif; CysL, cysteine loop; SH3 binding (1–3) Src homology 3 binding domains. (B) Color-coded peptide sequence of ADM-2 corresponding to panel A. For additional details see S2 Fig. (C) Percentage of viable (non-molting-defective) *nekl-2; nekl-3* adults in the indicated CRISPR-derived *adm-2* mutant backgrounds. Error bars represent 95% confidence intervals. p-Values were determined using Fisher’s exact test: ****p ≤ 0.0001; ns, not significant. (D–F) Predicted three-dimensional protein structures (for amino acid region 307–328) of wild-type ADM-2 (orange) superimposed on modeled structures for the Furin-1 (D; red), DM (E; cyan), and CysL (F; violet) mutant proteins. Three conserved histidine residues (His312, His316, His322) of the Zn-metalloprotease domain are represented as sticks. (F) A predicted 6.1-Å shift of the ‘tele’ nitrogen atom in the imidazole ring of His322 in the CysL mutant.

### The protease domain of ADM-2 is critical for its function

Most ADAM family members, including the human meltrins, contain a catalytically active metalloprotease domain that is distinguished by three conserved histidine residues (HExxHxxGxxH) that coordinate the binding of zinc [49, 50]. To determine if the putative metalloprotease activity of ADM-2 is critical for its influence on molting, we CRISPR-engineered an ADM-2 variant in which the three conserved histidine residues within the predicted Zn-binding domain were altered [Zn-MTP: H312–H322 (**HE**L**GH**TF**G**MD**H** > **DA**L**AY**TF**R**MD**Y**)] (Fig 5A and 5B). Notably, this variant displayed robust suppression of *nekl-2; nekl-3* molting defects, indicating that the protease function of ADM-2 is central to its function in molting and that its loss is sufficient for *nekl* suppression (Fig 5C).

To further assess ADM-2 functional domains, we tested CRISPR variants designed to disrupt a predicted N-terminal furin-cleavage site (NT-Furin: R149–R152 [**R**KK**R** > **V**KK**V**]) and a predicted disintegrin motif (DM: E388–G396 (EPGE**ECDCG** > EPGE**VLADP**) (Fig 5A and 5B). Both variants led to relatively weak suppression of *nekl-2; nekl-3*, suggesting that these domains are less critical with respect to the molting-associated functions of ADM-2 (Fig 5C). Interestingly, a variant that affects a predicted cysteine loop within the greater disintegrin domain (CysL: C438– P459 [CRAAIGICDL**DEYCNG**ETNDCP > CRAAIGICDL**QQNGDH**ETNDCP]) strongly suppressed *nekl-2; nekl-3* molting defects (Fig 5A–C). To better understand the basis for the observed suppression patterns, we made use of the AlphaFold database [51, 52] to obtain the predicted structure of ADM-2 and then performed homology modeling using the online Robetta protein structure prediction service [53, 54] to model the effects of our mutations. Notably, whereas the NT-furin and DM mutations were projected to have minimal impacts on the configuration of histidine residues within the Zn-MTP domain (Fig 5D and 5E), the CysL variant was predicted to substantially alter the position of His322, leading to an expected reduction in metalloprotease activity (Fig 5F). Collectively, these data indicate that the protease domain of ADM-2 is central to its impact on the molting process and that other N-terminal domains may play more minor or indirect roles in this context.

We also tested the importance of the ADM-2 C-terminal domain (R696–K952) using several CRISPR-generated lines. Deletion of most of the C terminus (CT: ΔH718–M942) led to strong suppression of *nekl-2; nekl-3* molting defects, whereas perturbation of individual SH3-binding domains led to suppressive effects ranging from minimal (SH3-2: P839–V853 [**P**NVQ**PPP**V**PRP**S**DDV** > **G**NVQ**GAG**V**GAG**S**LLE**] and SH3-3: K874–K884 [**K**TL**P**L**PPP**L**PK** > ITL**E**L**GAG**L**GL**]) to moderate (SH3-1: V722–P731 [**VP**V**RK**A**PPPP** > **EG**V**LA**A**GAVG**]) (Fig 5A–C). We also note the presence of a potential fourth SH3-binding domain in the region of P907 (Fig 5b, S2 Fig). As expected, none of the C-terminal mutations led to predicted changes in the configuration of histidine residues in the Zn-binding domain (S6 Fig). A role for the C-terminus in mediating ADAM function is not unexpected given that this domain is proposed to be important for the regulation of cell signaling and subcellular localization and contains specialized motifs that are thought to be involved in the ‘inside-out’ regulation of ADAM metalloprotease activity [30, 31, 33, 55].

### Overexpression of ADM-2 causes molting defects

Given that loss of ADM-2 did not lead to defects in molting but was capable of suppressing molting defects in *nekl-2; nekl-3* mutants, we hypothesized that ADM-2 may normally exert an inhibitory effect on the molting process. To test if ADM-2 is a negative regulator of molting, we generated strains carrying the *adm-2* cDNA under the control of a heat shock–inducible promoter (*P*_*hsp-16*_*::adm-*2 and *P*_*hsp-16*_*::adm-2::GFP*. Strikingly, when subjected to heat shock during larval development (i.e., overexpression of *adm-2*), we observed molting defects in ~50% of larvae carrying these heat-shock–ADM-2 transgenes (Fig 6A and 6D). In contrast, molting defects were not observed in heat-shocked controls or in non-heat-shocked worms (Fig 6A); ADM-2::GFP expression was specifically detected in heat-shocked worms only (Fig 6B and 6C). In addition, we observed a low frequency of blister phenotypes in animals that overexpressed ADM-2::GFP, a defect associated with detachment of the cuticle from the epidermis (Fig 6E). These findings are consistent with ADM-2 playing an inhibitory role in molting and suggest that ADM-2 may also influence attachment of the epidermis to the cuticle.

**Fig 6.**
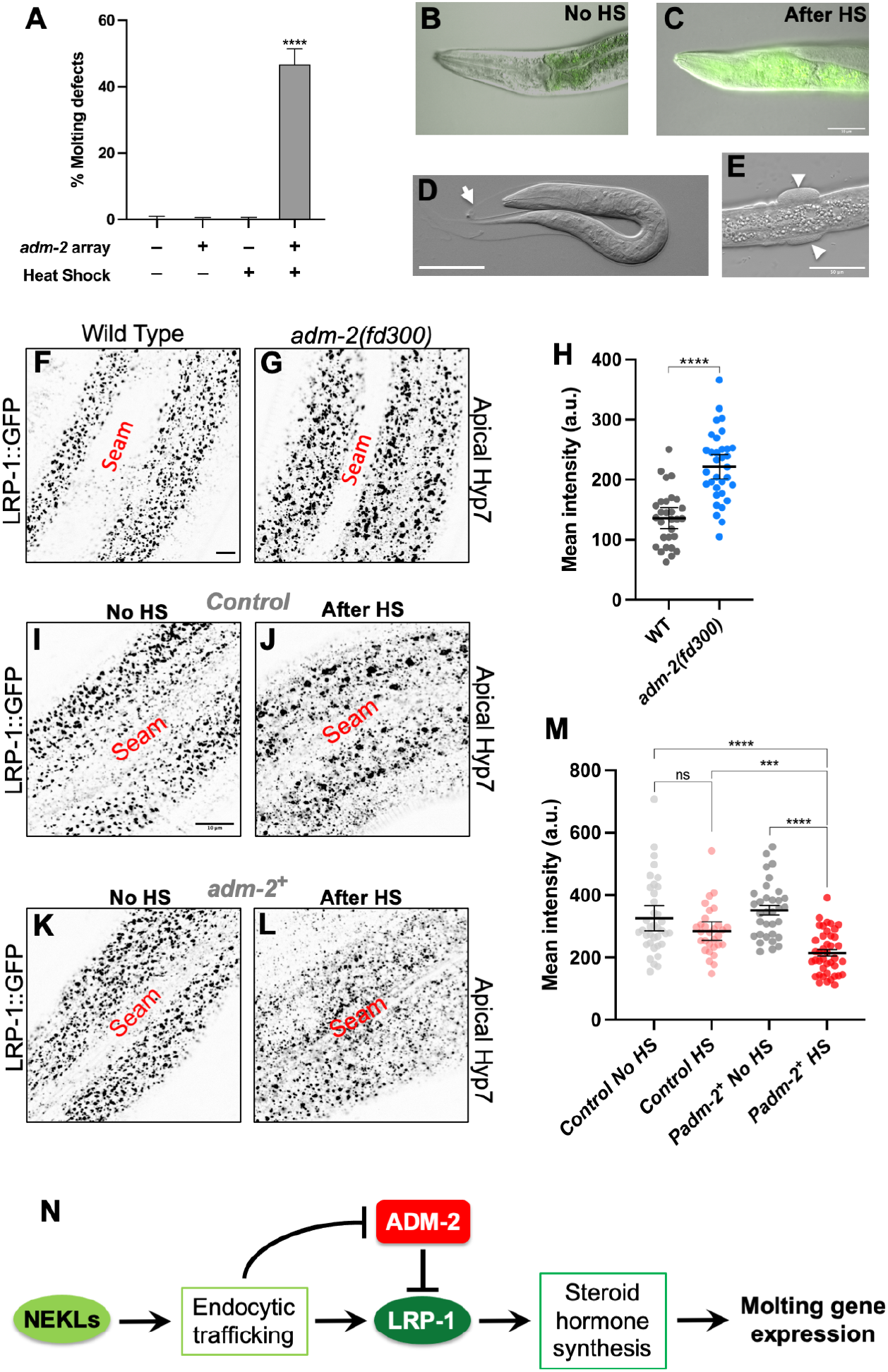
ADM-2 is a negative regulator of molting and LRP-1. Bar plot showing the percentage of molting defects in the specified strains in the presence and absence of the *P*_*hsp-16*_*::adm-2::GFP* transgene in the wild-type background and in the presence and absence of heat shock. Error bars represent 95% confidence intervals. p-Values were determined using Fisher’s exact test: ****p ≤ 0.0001. (B,C) Merged DIC and fluorescence images of an adult worm carrying the *P*_*hsp-16*_*::adm-2::GFP* transgene in the absence of heat shock (B) and after heat shock (C). (D) Molting-defective larva after heat shock. White arrow in D indicates unshed old cuticle. (E) Rare blister phenotype in a worm after ADM-2::GFP induction. White arrowheads in E indicate blisters. Bar in C = 50 μm (for B and C); in D = 50 μm; in E = 50 μm. (F,G) Representative confocal images of 1-day-old adult wild-type (F) and *adm-2(fd300)* (G) worms expressing LRP-1::GFP in the hyp7 region of the hypodermis. Bar in F = 5 μm (for F,G). (H) Dot plot showing LRP-1::GFP mean intensity (a.u.) within the apical plane of the worm hypodermis for each individual worm of the specified genotype. (I–L) Representative confocal images of 1-day-old adult worms expressing LRP-1::GFP in the hyp7 region of the hypodermis. (I,J) LRP-1 expression in wild-type control worms. (K,L) LRP-1 expression in worms containing the *P*_*hsp-16*_*::adm-2* transgene. Worms are shown in the absence of heat shock (I,K) and after heat shock (J,L). Bar in I = 10 μm (for I–L). (M) Dot plot showing LRP-1::GFP mean intensity (a.u.) within the apical plane of the worm hypodermis for each individual worm of the specified genotype and heat shock conditions. In H and M, group means along with 95% confidence intervals (error bars) are indicated. p-Values were obtained by comparing means using an unpaired t-test: ****p ≤ 0.0001; ***p ≤ 0.001; ns, p ≥ 0.05. (N) Genetic model of ADM-2 function in *C. elegans* molting.

### ADM-2 is a negative regulator of LRP-1/megalin

ADAM/meltrin family proteins function as “sheddases”, cleaving target peptides that are positioned at or near the outer leaflet of the plasma membrane [30, 31, 56–70]. Given our above findings, we hypothesized that ADM-2 may function as a sheddase for one or more apical membrane proteins that positively regulate the molting process. As such, loss of ADM-2 may lead to the upregulation or de-repression of these proteins, thereby alleviating molting defects in weak *nekl* loss-of-function mutants. One candidate protein for regulation by ADM-2 is LRP-1, a positive regulator of molting and a known target of NEKL trafficking (also see above) [71–79]. Notably, two mammalian homologs of LRP-1 (LRP1/LRP2) are negatively regulated by matrix metalloproteinases and ADAM family members, including the meltrin ADAM12 [71–81]. To test this possibility, we examined levels of LRP-1::GFP in *adm-2* null mutants and observed apical levels of LRP-1::GFP to be ~1.6-fold higher in *adm-2* mutants relative to wild type (Fig 6F–H). In contrast, loss of *adm-2* did not alter the levels of a clathrin heavy chain reporter at the apical membrane (S7A–D Fig), indicating that ADM-2 does not generally affect CME.

To further examine the relationship between ADM-2 and LRP-1 we tested the effects of ADM-2 overexpression on LRP-1::GFP levels. One-day-old adult worms carrying the *P*_*hsp-16*_*::adm-2* transgene and LRP-1::GFP were heat shocked, and LRP-1::GFP levels were assayed ~2–3 hours later. Whereas heat shock alone did not significantly alter LRP-1::GFP levels in the absence of the *P*_*HS*_*::adm-2* transgene, levels of LRP-1::GFP were ~1.6-fold lower when worms containing the *P*_*HS*_*::adm-2* transgene were heat shocked relative to non-heat-shocked controls (Fig 6I–M). These striking reciprocal findings are consistent with ADM-2 functioning as a negative regulator of LRP-1 and suggest that suppression by loss of functional ADM-2 may occur in part through the upregulation or de-repression of cargo including LRP-1.

## Discussion

In this study we demonstrated a unique role for the sole *C. elegans* meltrin family member, ADM-2, as a negative regulator of the molting process. Whereas loss of ADM-2 did not overtly impact the molting cycle, overexpression of ADM-2 during larval development did lead to molting defects and larval arrest. Moreover, our previously described non-biased forward genetic screen identified two independent loss-of-function alleles of *adm-2* as suppressors of larval arrest and molting defects in *nekl-2; nekl-3* mutants [29]. Previous studies have implicated proteases as positive regulators of molting including NAS-36, NAS-37, CPZ-1, and SURO-1 [82–87]. Extracellular proteases have been suggested to be important for the detachment and degradation of the old cuticle and may also play a dynamic role in ECM remodeling during new cuticle synthesis [2]. Interestingly, roles for protease inhibitors in molting have also been reported, as loss of the Kunitz domain–containing protease inhibitors MLT-11 and BLI-5 lead to molting defects [83, 88, 89]. Protease inhibitors have been suggested to be important for temporally or spatially restricting the activity of extracellular proteases during the molting cycle. Consistent with this, epidermally expressed proteases and protease inhibitors are highly enriched among genes that are transcriptionally regulated with the molting cycle, indicating tight control of their proteolytic activity [90, 91].

Although we previously reported the suppression of *nekl* molting defects by mutations affecting genes closely connected to CME, our data do not support a role for ADM-2 in the regulation of intracellular trafficking per se. Rather, our findings are consistent with ADM-2 being a cargo of CME, including during its passage through endosomes and its turnover in lysosomes. Moreover, we observed increased levels of ADM-2 after NEKL knockdown along with the retention of ADM-2 in intracellular compartments of the epidermis. Collectively these findings suggest that loss of NEKL functions may lead to increased or aberrant ADM-2 activity, which may contribute to molting defects in these mutants.

Notably, we identified LRP-1 as a candidate target of ADM-2 sheddase activity. Epidermal LRP-1 levels were increased in *adm-2* null mutants and were reduced when ADM-2 was overexpressed. Moreover, alteration of the ADM-2 metalloprotease domain was sufficient to mediate strong suppression of *nekl* molting defects, indicating that proteolytic/sheddase activity is central to the effects of ADM-2 on molting. Notably, human LRP1 is a known target for cleavage by ADAM10, ADAM12, and ADAM17 [71–79]. Moreover, after cleavage by ADAMs, the solubilized extracellular domain of LRP1 retains its ability to bind apolipoproteins with high affinity. This was proposed to decrease the effective concentration of ligand available for internalization by membrane-bound LRP1, leading to a reduction in cholesterol internalization [80]. In addition, matrix metalloproteases have been proposed to mediate the cleavage and proteolysis of mammalian LRP2/megalin, although the identity of the protease(s) was not determined [77–81]. Given that *C. elegans* LRP-1 is most similar to mammalian LRP2, our study suggests that LRP2 may be regulated by meltrin family members. Together, our data are consistent with ADM-2 playing a conserved role in the repression of LRP family members.

Our findings, together with published data from our lab and other research groups, point to a working model in which epidermal intracellular trafficking may play two roles in the molting process (Fig 6N). Based on previous studies, endocytosis is required for the internalization of factors required for molting, such as steroid hormone precursors [5, 6], and may also be important for recycling old cuticle components, such as collagens [47, 92]. This function is consistent with our recent observation that loss of *nekls* leads to defects in the transcriptional upregulation of those molting genes for which their expression depends on the activation of nuclear hormone receptors [12]. In addition, endocytosis may also be required for the uptake and degradation of cargo that would otherwise exert an inhibitory effect on molting. Impaired endocytic trafficking in *nekl* mutants may simultaneously lead to a reduction in cholesterol uptake via LRP-1 and an increase in the levels of a negative regulator of LRP-1, ADM-2 (Fig 6N). In this model, loss of *adm-2* would lead to de-repression of LRP-1, and possibly other positive effectors of molting, thereby compensating for a partial loss of NEKL trafficking functions. In contrast, when NEKL functions are severely reduced, loss of ADM-2 would not be expected to offset a more severe deficiency in the uptake of steroid precursors, consistent with our observation that *adm-2* mutations do not suppress strong loss-of-function alleles in *nekls*. In summary, our findings expand the roles for NEKLs and intracellular trafficking in the molting process and implicate ADM-2 as a negative regulator of the molting process.

## Materials and Methods

### Strains

*C. elegans* strains were maintained according to standard protocols [93] and were propagated at 22°C, unless stated otherwise. Strains used in this study are listed in S1 Table.

### Transgenic Rescue

Fosmids containing rescuing sequences for *adm-2*/C04A11.4 (WRM0620dD12, WRM0632aG02, and WRM0610cA04, 2–6 ng/μl each + *sur-5::RFP* [pTG96], 50–100 ng/μl) were injected into strain WY1342. Stable strains (WY1386 and WY1388) containing rescuing arrays for both *nekl-3* (*fdEx286*; GFP^+^) and *adm-2* (*fdEx315* or *fdEx356*; RFP^+^) were scored to determine the percentage of viable RFP^+^ progeny.

### Determination of dominant versus recessive alleles

To distinguish between dominant and recessive alleles, we first crossed suppressed *nekl-2*(*fd81*); *nekl-3*(*gk894345) fd130* hermaphrodites to WY1145 [*nekl-2*(*fd81*); *nekl-3*(*gk894345); fdEx286 (nekl-3^+^ + sur-5::GFP)*] males and scored for suppression of GFP^−^ cross-progeny males. For *fd130* and *fd162*, 50/96 and 30/52 viable cross-progeny adult males were GFP^−^, respectively, indicating that these mutations are either dominant or on LGX. We next crossed *nekl-2*(*fd81*); *nekl-3*(*gk894345) fd130* hermaphrodites to WY1232 [*nekl-2*(*fd81*); *nekl-3*(*gk894345*); *fdEx186* (*nekl-3*^*+*^ + *sur-5::GFP*); *fdEx197 (sur-5::RFP)*] males and scored for suppression of GFP^−^ RFP^+^ cross-progeny hermaphrodites. In the case of cross-progeny *fd130/+* adult hermaphrodites, 62/62 were either GFP^+^ RFP^−^ or GFP^+^ RFP^+^; no GFP^−^ RFP^+^ adults were observed. Similarly, 182/185 *fd162/+* adult hermaphrodites were either GFP^+^ RFP^−^ or GFP^+^ RFP^+^, and only 3/185 were GFP^−^ RFP^+^. Given the ~98.5% penetrance of *nekl-2*(*fd81*); *nekl-3*(*gk894345)* larval lethality [29], our results indicate that *fd130* and fd162 are fully recessive but are on LGX.

### RNAi

Primers containing the binding motif for T7 RNA polymerase (5’-TAATACGACTCACTATAGGGAGA-3’) and corresponding to *adm-2* (5’-GACCACAACAATGATACGGTCGAA-3’; 5’-CCTGGACACAATGCAGCATTTTGA-3’), *unc-71* (5’-TGTCGTCGACGGTTCCGAAGA-3’; 5’-GCATCAGACAGACCAGGCATAG-3’), *adm-4* (5’-ATGCATTCAATACACGTGTGA-3’; 5’-CTTCCTCTCCCAGATATATCGT-3’), *sup-17* (5’-AGTGTCAACCTGGTCTTCCTG-3’; 5’-CTGTGCCCATTGTGTTAGAGTTTC-3’), *mig-17* (5’-CTCAGCTACACAAGGAATGGC-3’; 5’-TTCGCACACGTTCTACAACA-3’), *tag-275* (5’-TGTTCTCGCGTCATTCGTTGC-3’; 5’-ACTCGGTTTATTGGAACATTTGGC-3’), *F27D9.7 (*5’-CAACATTCTGTGCGATGCGGT-3’; 5’-TTAAATGGGCGCGACAGATCC-3’), *adt-1* (5’-GTCAGTGCACTCACTGGACAT-3’; 5’-GGTTAGGCATGGCCTGAATCT-3’), *adt-2* (5’-GAAGACGAAACCGAAGTCTGC-3’; 5’-TTACCTCCCCATGCAGCATTT-3’), *adt-3* (5’-CAGGTATGTAACGGTGACTCCA-3’; 5’-CATTACACATGGTCCGGTTTC-3’), and *gon-1* (5’-TGGATCACTGAAGATGTGTCT-3’; 5’-GCACTCCAATCAGTATTTCTC-3’) were used to generate dsRNA using standard methods [94]. After injection at 0.8–1.0 μg/μl into WY1145 hermaphrodites, F1 progeny were scored for adult viability. For RNAi feeding experiments, the relevant bacterial strains were obtained from Geneservice and IPTG (8 mM) was added to growth plates [95]. Worm strains were grown on *lin-35(RNAi)* plates for two generations to increase RNAi susceptibility [96]. Second-generation fourth larval stage (L4) worms growing on *lin-35(RNAi)* plates were transferred to experimental plates and were imaged after 48 hours. RNAi feeding experiments were performed at 20°C.

### ADM-2 CRISPR mutant alleles

*D*esign of repair sequences containing introduced restriction sites was facilitated using CRISPRcruncher [97]. For details on primers sequences see S2 File.

#### Strong loss-of-function alleles

Alleles *fd228–fd230* and *fd235–fd237* were generated using guide dual sequences SB1 and SB2, PCR amplification primers SB3 and SB4, and the sequencing primer SB6. In *fd228–fd230* and *fd235–fd237*, an ~3.2-kb region spanning *adm-2a* exons 3–9 is deleted. *fd228* is an indel predicted to encode sequences through T122 of ADM-2a, followed by eight divergent amino acids and a stop codon. *fd229* is an indel predicted to encode sequences through S123 of ADM-2a, followed by a stop codon. *fd230* is an indel predicted to encode sequences through T122 of ADM-2a, followed by a stop codon. *fd235* is an indel predicted to encode sequences through T122 of ADM-2a, followed by 23 new amino acids and a stop codon. *fd236* is an indel predicted to encode sequences through F118 of ADM-2a, followed by six new amino acids and a stop codon. *fd237* is an indel predicted to encode sequences through T122 of ADM-2a, followed by a single new amino acid and a stop codon. Alleles *fd298–302* were generated using guide sequences SB46 and SB47, PCR amplification primers SB48–SB51, and the sequencing primer SB52. In *fd298–fd302*, *fd313*, *fd316*, and *fd317* an ~7.4-kb region spanning *adm-2a* exons 1–19 is deleted. *fd298* is an indel that encodes sequences through T2 of ADM-2a followed by four new amino acids and a stop codon. *fd299* is a deletion that encodes sequences through M1 of ADM-2A, followed by three new amino acids and a stop codon. *fd300* is an indel that encodes sequences through D3 of ADM-2a, followed by 14 new amino acids and a stop codon. *fd301* and *fd302* are identical indels that encode sequences through D3 of ADM-2a, followed by 12 new amino acids and a stop codon. *fd313* encodes sequences through T2 of ADM-2a, followed by 40 new amino acids and a stop codon. *fd316* deletes the normal start codon; an alternative ATG is predicted to encode 24 new amino acids followed by a stop codon.

*fd317* encodes sequences through D3 of ADM-2a, followed by 16 new amino acids and a stop codon. *fd318* encodes sequences through D3 of ADM-2a, followed by 8 new amino acids and a stop codon.

#### *adm-2* metalloprotease mutation (Zn-binding domain)

Alleles *fd243–fd247* were generated using the guide sequence SB27, the repair template Rep2, PCR amplification primers SB28 and SB29, and sequencing primers SB31 and SB32. *fd243–fd247* change the predicted ADM-2a Zn-metalloprotease active site spanning H312–H322 (**HE**L**GH**TF**G**MD**H**) to **DA**L**AY**TF**R**MD**Y**(altered aa are in bold). The Zn-metalloprotease consensus motif is HEXXHXUGUXH, where U is an amino acid containing a bulky hydrophobic residue. The edited locus contains an introduced BstBI site.

#### *adm-2* disintegrin motif mutation

Allele *fd322* was generated using guide sequence DF1, the repair template Rep9, PCR amplification primers DF2 and DF3, and sequencing primers DF4 and DF5. *fd322* changes the predicted ADM-2a disintegrin motif (DM) spanning E388–G396 (EPGE**ECDCG**) to EPGE**VLADP**. The edited locus contains introduced NheI and BamHI sites.

#### *adm-2* cysteine loop mutation

Alleles *fd324–fd325* were generated using guide sequence DF6, the repair template Rep10, PCR amplification primers DF2 and DF3, and sequencing primers DF4 and DF5. *fd324–fd325* change the predicted ADM-2a cysteine loop (CysL) spanning C438–P459 (CRAAIGICDL**DEYCNG**ETNDCP) to CRAAIGICDL**QQNGDH**ETNDCP. The edited locus contains an introduced PstI site.

#### *adm-2* furin 1 mutation

Alleles *fd288-fd290* were generated using guide sequence SB17, the repair template Rep6, PCR amplification primers SB18 and SB19, and sequencing primers SB20 and SB21. *fd288-fd290* change the predicted ADM-2a furin-1 cleavage site spanning R149–R152 (**R**KK**R**) to **V**KK**V**. The edited locus contains an introduced BamHI site.

#### *adm-2* SH3-binding domain 1 mutation

Alleles *fd248–fd251* were generated using the guide sequence SB32, the repair template Rep3, PCR amplification primers SB16 and SB34, and sequencing primer SB35. *fd248–fd251* alter the predicted ADM-2a SH3-1 domain spanning V722–P731 (**VP**V**RK**A**PPPP)**to **EG**V**LA**A**GAVG**. The edited locus contains an introduced XhoI site. *adm-2* SH3-binding domain 2 mutation. Alleles *fd252–fd256* were generated using the guide SB36, the repair template Rep4, PCR amplification primers SB37 and SB38, and sequencing primers SB39 and SB40. *fd252–fd256* alter the predicted ADM-2a SH3-binding domain 2 domain spanning P839–V853 (**P**NVQ**PPP**V**PRP**S**DDV**) to **G**NVQ**GAG**V**GAG**S**LLE**. The edited locus contains an introduced XhoI site.

#### *adm-2* SH3-binding domain 3 mutation

Alleles *fd257–fd259* were generated using the guide sequence SB41, the repair template Rep5, PCR amplification primers SB37 and SB38, and sequencing primers SB39 and SB40. *fd257–fd259* alter the predicted ADM-2a SH3-binding domain 3 spanning K874–K884 (**K**TL**P**L**PPP**L**PK**) to **I**TL**E**L**GAG**L**GL**. The edited locus contains an introduced XhoI site.

#### *adm-2* C-terminal (cytoplasmic domain) deletion

Alleles *fd231–fd234* were generated using the guide sequences SB12 and SB13, the repair template Rep1, PCR amplification primers SB14 and SB15, and sequencing primer SB16. *fd231–fd234* remove sequences from H718 to M942 of ADM-2a and introduce a XhoI site.

### ADM-2 expression plasmids and strains

*adm-2::GFP* and *adm-2::mScarlet* endogenously tagged strains were made using CRISPR/Cas9 technology in collaboration with SunyBiotech Corporation (China). Expression vectors were generated by amplifying the *adm-2* promoter region from fosmid WRM0620dD12 using primers SB57 and SB58. After digestion with SphI and SalI, the ~2.1-kb PCR product was inserted into pPD95.75 to create pDF403–pDF405. *adm-2* cDNA was amplified from plasmid pDONR201 (Horizon Inc.) using primers SB59 and SB60. Digestion and ligation of the ~2.8-kb PCR product and pDF403 with XmaI and KpnI generated pDF417–pDF420, which were confirmed by sequencing. pDF417 (~100 ng/μl) was injected into N2 worms with *sur-5::RFP* (~50 ng/μl) to obtain lines carrying extrachromosomal arrays (*fdEx353*, *fdEx354*). CRISPR-tagged *adm-2::mScarlet* and *adm-2::GFP* strains contain codon optimized fluorescent-reporter insertions just preceding the *adm-2a* stop codon (SUNY biotech). In addition, sequences upstream of the stop codon contain the indicated (bold) silent mutations (LGX 13695343–13695387).

TC**T**GAAGATGCAGCTGCAACCGAAGAAAAAGTAGATGTTCGCTC**C**(wild type)

TC**A**GAAGATGCAGCTGCAACCGAAGAAAAAGTAGATGTTCGCTC**G**(CRISPR tagged)

### *adm-2* and *adm-2::GFP* heat shock strains

An ~4.6-kb *adm-2::GFP::unc-54* 3’UTR cDNA product was obtained by digesting pDF420 with XbaI and ApaI enzymes. Digestion of heat shock vectors pPD49.78 and pPD49.83 was performed using NheI and ApaI to obtain an ~3-kb vector backbone. Ligation of the *adm-2* cDNA with vector backbones pPD49.83 and pPD49.78 generated pDF429–pDF431 and pDF432–pDF434, respectively. Likewise, an ~2.8-kb *adm-2::GFP* product was obtained by digesting pDF420 with XbaI and KpnI enzymes. Digestion of heat shock vectors pPD49.78 and pPD49.83 was performed using NheI and KpnI to obtain an ~3-kb vector backbone. Ligation of *adm-2::GFP* with vector backbones pPD49.83 and pPD49.78 generated pDF423–pDF425 and pDF426–pDF428, respectively. pDF430 and pDF433 (50 ng/μl each) were injected into N2 with pRF4 [*rol-6(gf)*] (~50 ng/μl) to obtain lines carrying extrachromosomal arrays (N2: *fdEx373–375*). Likewise, pDF424 and pDF425 (50 ng/μl each) were injected into N2 with pRF4 [*rol-6(gf)*] (~50 ng/μl) to obtain lines carrying extrachromosomal arrays (*fdEx381, fdEx382*).

### Heat shock methods

For Fig 6A, all worm strains were synchronized using bleach. Synchronized L1 worms were plated on NGM plates and grown at 20°C for 2 hours. The plates were then incubated at 34°C for 4 hours, after which they were shifted back to 20°C for 20 hours. The plates were shifted again to 34°C for 4 hours and were subsequently grown at 20°C for 20–24 hours before the percentage of molting-defective worms was determined. For Fig 6I–L, 1-day-old adult worms grown at 20°C were heat shocked at 34°C for 4 hours and were shifted to 20°C for 2–3 hours before imaging.

### Protein domain identification and alignment tools

The following sites were used to identify domains within ADM-2 and human ADAM homologs:

http://nls-mapper.iab.keio.ac.jp/cgi-bin/NLS_Mapper_form.cgi

https://www.ncbi.nlm.nih.gov/Structure/cdd/wrpsb.cgi

https://www.ebi.ac.uk/interpro/

https://prosite.expasy.org/

https://psort.hgc.jp/form2.html

http://www.cbs.dtu.dk/services/TMHMM/

http://www.cbs.dtu.dk/services/SignalP/

http://phobius.sbc.su.se/

http://tcoffee.crg.cat/apps/tcoffee/do:regular

https://embnet.vital-it.ch/software/BOX_form.html

### Image acquisition

Fluorescence images in the following figure panels—Fig 2A–C; Fig 6F–G; S3G Fig; and S7 Fig—were acquired using an Olympus IX81 inverted microscope with a Yokogawa spinning-disc confocal head (CSU-X1). Excitation wavelengths were controlled using an acousto-optical tunable filter (ILE4; Spectral Applied Research). MetaMorph 7.7 software (MetaMorph Inc.) was used for image acquisition. z-Stack images were acquired using a 100×, 1.40 N.A. oil objective. Fig 3A–G; Fig 3A’–G’; Fig 4A–L; Fig 6I–L; S3A–F Fig; S3A’–F’ Fig; and S4C–H Fig were acquired using an Olympus IX83 inverted microscope with a Yokogawa spinning-disc confocal head (CSU-W1). z-Stack images were acquired using a 100×, 1.40 N.A. silicone oil objective. cellSense3.1 software (Olympus corporation) was used for image acquisition. DIC images in the panels Fig 1A–C and Fig 7B–E were acquired using a Nikon Eclipse epiflourescence microscope and the cellSense3.1 software (Olympus corporation).

### Image analysis

Mean intensity (measured in arbitary units, a.u.), percent of fluorescence-positive pixels above threshold, and the colocalization analysis were performed using Fiji software (NIH; available at https://imagej.net/Fiji/Downloads). For a given z-plane of interest, rolling ball background subtraction was performed (radius = 50 pixels), and the polygon selection tool was used to choose the region of hyp7 in which the mean intensity was quantified (Fig 4, Fig 6, S5 Fig, and S7 Fig). The percentage of fluorescence-positive pixels for the region of interest (Fig 2) was determined after thresholding, and the “Huang” thresholding algorithm was used for strain comparisons. For colocalization, rolling ball background subtraction was performed (radius = 25 pixels), followed by use of the mean filter (radius = 2 pixels) to minimize noise. Finally, the same thresholding algorithm was used for one particular channel to obtain binary images to be used as masks (GFP::CHC-1 and HGRS-1::GFP—“Otsu”; ADM-2:mScarlet—“Isodata”). This binary mask was combined using the “AND” boolean operation to the original image and the combined image was used for the colocalization analysis. Data from Fig 6F and 6G (Olympus IX81) and Fig 6I–L (Olympus IX83) were obtained on different confocal microscopes and thus differ in their mean intensity values.

### Auxin treatment

Auxin (indole-3-acetic acid) was purchased from Alfa Aesar. A 100× stock auxin solution (0.4 M) was made by dissolving 0.7 g of auxin in 10 ml of 100% ethanol. A mixture of 25 μl of stock auxin solution and 225 μl of distilled water was added to plates containing 1-day-old adult worms.

### Protein 3D structure analysis

PDB file (Identifier : AF-G5EDW5-F) containing the three-dimensional structure details of *C. elegans* ADM-2 was obtained from the AlphaFold database ( https://alphafold.ebi.ac.uk/) [51, 52]. Using the AlphaFold structure as template, homology modeling was performed by the online Robetta structure prediction server (https://robetta.bakerlab.org/) to obtain the predicted the three-dimensional structures of the respective ADM-2 mutants [53, 54]. For modeling CM (Comparative modeling) option was used and the number of models to sample was selected as 1. Other options remained unchanged. The homology modeled three-dimensional structures were rendered, and was superimposed onto the AlphaFold structure of ADM-2 using the PyMOL 2 software (The PyMOL Molecular Graphics System, Version 2.0 Schrödinger, LLC.).

### Statistical analyses

Statistical analyses were carried out using Prism software (GraphPad) following established standards [98].

## Supporting information

Combined Supplementary Figures

S1 Table. List of strains

S1 File. Complete raw data

S2 File. List of primers

## Acknowledgements

We thank Amy Fluet for editing this manuscript and Barth D. Grant for the HGRS-1 marker strain. This project was supported by R35 GM136236 to DSF and by an Institutional Development Award (IDeA) from the National Institute of General Medical Sciences of the National Institutes of Health (P20GM103432).

---

**S1 Fig. Loss of other *C. elegans* ADAM family members does not suppress *nekl* defects**

(A) Dot plot showing average brood sizes for 10 individual wild-type and *adm-2(fd300)* mutant worms. (B) Bar plot showing the failure of most *C. elegans* ADAM family members to suppress molting defects in *nekl-2; nekl-3* mutants. (C) Table showing ADM-2 *C. elegans* orthologs and their corresponding human homologs. Error bars in A,B represent 95% confidence intervals. p-Values were determined using an unpaired t-test (A) (ns, p ≥ 0.05) or Fisher’s exact test (B): ****p ≤ 0.0001.

**S2 Fig. Alignment of *C. elegans* ADM-2 with human ADAMs**

Peptide alignment of *C. elegans* ADM-2 with human meltrin family members (ADAM9/12/19/33). Predicted domains of ADM-2 are color coded. NLS, nuclear localization domain.

**S3 Fig. Additional ADM-2 expression**

(A–F and A’–F’) Representative DIC (A–F) and confocal (A’–F’) images of ADM-2 expression showing the anterior hypodermis (A, A’), nerve ring (B, B’), tail neurons (C, C’), and various stages of embryonic development (D–F’). Bar in A’ = 10 μm (for A, A’); in B’ = 10 μm (for B, B’); in C’ = 10 μm (for C, C’); in F’ = 10 μm (for D–F’). (G) Representative confocal image of an L2 larva expressing multi-copy P_*adm-2*_::ADM-2::GFP in the plasma membrane of head neurons. Bar in G = 25 μm.

**S4 Fig. Supplemental colocalization data**

(A,B) Dot plots showing quantification of Mander’s overlap coefficient for the overlap of ADM-2::mScarlet with GFP::CHC-1 and P_hyp7_::HGRS-1::GFP proteins within the apical (A) and medial planes. Mean values and 95% confidence intervals (error bars) are indicated. p-Values were calculated using an unpaired test: ****p ≤ 0.0001, **p ≤ 0.01. (C–H) Representative confocal images of GFP::CHC-1 (C), P_hyp7_::HGRS-1::GFP (F), and ADM-2::mScarlet (D,G) within the hyp7 medial plane. C’–H’ are insets of C–H confocal images. Bar in E= 10 μm (for C–H); in E’ = 5 μm (for insets C’–H’). (E’,H’) White arrows show ADM-2 large vesicular structures that do not colocalize with GFP::CHC-1 and P_hyp7_HGRS-1::GFP puncta, which are lysosomes. Cyan arrows indicate vesicles containing ADM-2 that colocalize with GFP::CHC-1 and P_hyp7_::HGRS-1::GFP.

**S5 Fig. ADM-2 levels are increased slightly upon weak loss of *mlt-3***

(A,B) Dot plot showing the mean intensity (a.u.) of GFP::CHC-1 (A) and ADM-2::mScarlet (B) expression in the presence of Control RNAi (i.e., empty vector) and *mlt-3* RNAi. Group means along with 95% confidence intervals (error bars) are indicated. p-Values were obtained by comparing means using an unpaired t-test: **p ≤ 0.01, *p ≤ 0.05.

**S6 Fig. Mutations in the C-terminal region of ADM-2 do not cause conformational changes to the metalloprotease active site**

Three-dimensional protein structure of the region including amino acids 307–328 of wild-type ADM-2 (orange) superimposed on modeled structures of SH3-1, SH3-2, and SH3-3 mutant proteins.

**S7 Fig. Clathrin expression is not perturbed by loss of ADM-2 function**

(A,B) Representative confocal images of GFP::CHC-1 expression in the hyp7 region of the hypodermis in the apical plane in wild-type (A) and *adm-2(fd300)* null mutant (B) 1-day-old adult worms. Bar in A = 10 μm (for A,B). (C,D) Dot plots showing GFP::CHC-1 mean intensity (a.u.) (C) and the percentage of GFP-positive pixels (D) within the apical plane for each individual worm of the specified genotype. In C and D, group means along with 95% confidence intervals (error bars) are indicated. p-Values were obtained by comparing means using an unpaired t-test. ns, p > 0.05.

**S1 Table. List of all the strains used in this study (pdf).**

**S1 File. Compilation or raw data used in this study (MS Excel).**

**S2 File. List of primers used in this study (MS Excel).**

