## Supplementary material for "An unexpected role for the conserved ADAM-family metalloprotease ADM-2 in *Caenorhabditis elegans* molting": Combined Supplementary Figures

**S1 Fig**

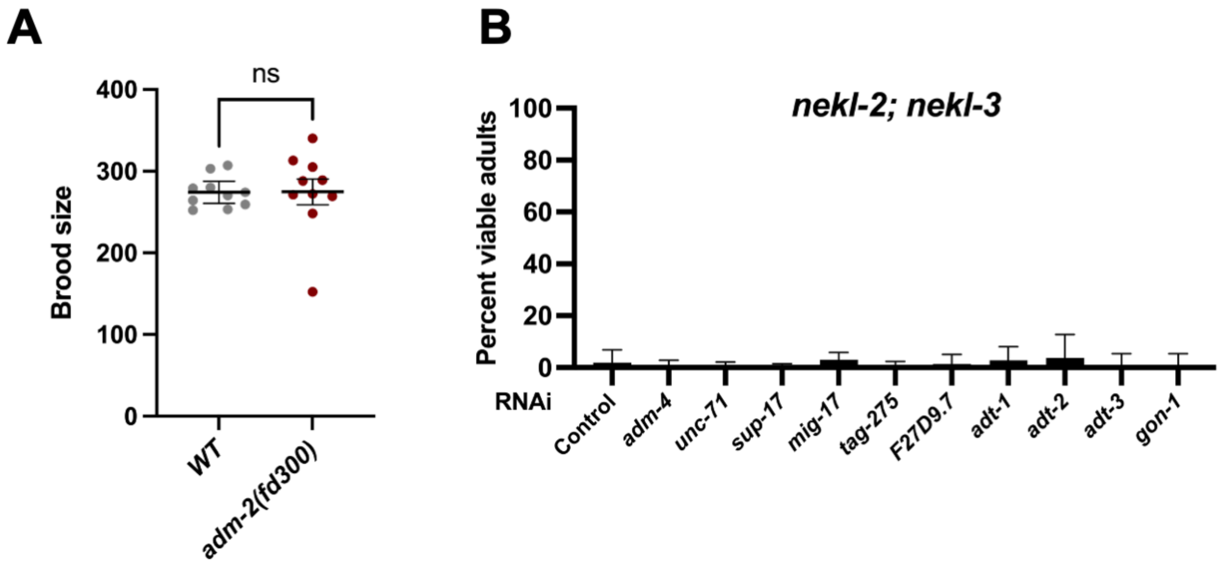


**C**

| *C. elegans* ADAMs | Human homologs |
|---|---|
| ADM-2 | ADAM9/12/19/33 |
| ADM-4 | ADAM17 |
| UNC-71 | ADAM15 |
| SUP-17 | ADAM10 |
| MIG-17 | ADAMTS6/16/18 |
| TAG-275 | ADAM28/ADAM DEC1 |
| F27D9.7 | ADAM28/ADAM DEC1 |
| ADT-1 | ADAMTS6/ADAMTS18 |
| ADT-2 | ADAMTS6/ADAMTS18 |
| ADT-3 | ADAMTS10/ADAMTS16 |
| GON-1 | ADAMTS20/ADAMTS9 |

**S1 Fig. Loss of other *C. elegans* ADAM family members does not suppress *nekl* defects**
(A) Dot plot showing average brood sizes for 10 individual wild-type and *adm-2(fd300)* mutant worms. (B) Bar plot showing the failure of most *C. elegans* ADAM family members to suppress molting defects in *nekl-2; nekl-3* mutants. (C) Table showing ADM-2 *C. elegans* orthologs and their corresponding human homologs. Error bars in A,B represent 95% confidence intervals. p-Values were determined using an unpaired t-test (A) (ns, $p \geq 0.05$) or Fisher's exact test (B): ****$p \leq 0.0001$.

### S2 Fig

```
ADM-2      1 MTDTLDL------------------------------------------KLSSR----RQ
ADAM9      1 MGSGARFPSGTLRVRWLLLLGLVGPVLG---AAR-PGFQQTS-------HLSSYEII-TP
ADAM12     1 MAAR-PLPVSPA--RALL-LALAGALLAPCEARGVSLWNQGRADEVVSASVGSG----DL
ADAM19     1 MPGG----AGAA--RLCL-LAFALQPLRPRAARE-PGWTRGSE--EGSPKLQHE-LIIPQ
ADAM33     1 ------------------------------------------------------------

ADM-2     15 WNPVRCPVRLEVDGSAQTPTSL--VQTALNNPSFDLVIAAPNGQNVYI-PFTEDRKLFTA
ADAM9     49 WRLTRERRE------APRP-YSKQVSYVIQAEGKEHIIHLERNKDLLPEDFVVYTYNKEG
ADAM12    53 WIPVKSFDS------KNHP-EV--LNIRLQRESKELIINLERNEGLIASSFTETHYLQDG
ADAM19    50 WKTSESPVR------EKHP-LK--AELRVMAEGRELILDLEKNEQLFAPSYTETHYTSSG
ADAM33     1 ----------------SKP-DM--GLVALEAEGQELLLELEKNHRLLAPGYIETHYGPDG

ADM-2     72 NIADDPSTS---SLISHCHFEGVTEDG-RHALSLCDPGEITGLLMTQTNRFGLSTSNNGS
ADAM9    102 --TLITDHP---NIQNHCHYRGYVEGVHNSSIALSDCFGLRGLLHL----------ENAS
ADAM12   104 --TDVSLARNYTVILGHCYYHGHVRGYSDSAVSLSTCSGLRGLIVF----------ENES
ADAM19   101 --NPQTTTR---KLEDHCFYHGTVRETELSSVTLSTCRGIRGLITVS---------SNLS
ADAM33    42 --QPVVLAP---NHTDHCHYQGRVRGFPDSWVVLCTCSGMSGLITLS---------RNAS
                                                    Cysteine switch
                                  Furin-1(R149-R152)
ADM-2    128 FVLIPYVEN--NCDLGSLVHSSSRKKRQIGKQ----------------NT-VIDRNPS--
ADAM9    147 YGIEPLQNS--S-H-FE--HIIYRMDDVYKEPLKCGVSNKDIEKETAKDEEEEPPSMTQL
ADAM12   152 YVLEPMKSA--T---NR--YKLFPAKKLKSVRGSCGSHHNTP-NLAAKNV-FPPPSQTWA
ADAM19   147 YVIEPLPDS--K---GQ--HLIYRSEHLKPPPGNCGFEHSKPTTRDWALQ-FTQQTKKRP
ADAM33    88 YYLRPWPPRGSKDF-ST--HEIFRMEQLLTWKGTCG--HRDPGNKAGMTS-LPGGPQS--

                     Protease domain(R177-P373)
ADM-2    167 -YIREHLDGRKRFVELALVADYSVYTKYDSDEKKVNDYMQQTMNILNSLYFPLNIRITLV
ADAM9    201 LRRRRAVLPQTRYVELFIVVDKERYDMMGRNQTAVREEMILLANYLDSMYIMLNIRIVLV
ADAM12   203 RRHKRETLKATKYVELVIVADNREFQRQGKDLEKVKQRLIEIANHVDKFYRPLNIRIVLV
ADAM19   199 RRMKREDLNSMKYVELYLVADYLEFQKNRRDQDATKHKLIEIANYVDKFYRSLNIRIALV
ADAM33   140 -RGRREARRTRKYLELYIVADHTLFLTRHRNLNHTKQRLLEVANYVDQLLRTLDIQVALT

ADM-2    226 HSEIWKKGDQISVIPDSKETLNNFMEYKKI-MLKDHFFDTGYLMTTLKFDEGVVGKAYKG
ADAM9    261 GLEIWTNGNLINIVGGAGDVLGNFVQWREKFLITRRRHDSAQLVLKKGFG-GTAGMAFVG
ADAM12   263 GVEVWNDMDKCSVSQDPFTSLHEFLDWRKMKLLPRKSHDNAQLVSGVYFQGTTIGMAPIM
ADAM19   259 GLEVWTHGNMCEVSENPYSTLWSFLSWRRK-LLAQKYHDNAQLITGMSFHGTTIGLAPLM
ADAM33   199 GLEVWTERDRSRVTQDANATLWAFLQWRRG-LWAQRPHDSAQLLTGRAFQGATVGLAPVE

                                       Zn binding(H312-H322)
ADM-2    285 TMCSYDYSGGIYVDHNNDTVETVATFAHELGHTFGMDHDPND-KDVCYCP--MPRCIMNP
ADAM9    320 TVCSRSHAGGINVFGQITVETFASIVAHELGHNLGMNHDDGR-D--CSCG--AKSCIMNS
ADAM12   323 SMCTADQSGGIVMDHSDNPLGAAVTLAHELGHNFGMNHDTLDRGCSCQMAVEKGGCIMNA
ADAM19   318 AMCSVYQSGGVNMDHSENAIGVAATMAHEMGHNFGMTHDSAD-C--CSASAADGGCIMAA
ADAM33   258 GMCRAESSGGVSTDHSELPIGAAATMAHEIGHSLGLSHDPDG-C-CVEAAAESGGCVMAA
                                               Disintegrin domain(A380-A467)
                                                 Disintegrin motif(E388-G396)
ADM-2    342 QSGH--MEVWSECSVKNLASGFNRGIDLCLFNEPGKK--PSDAKCGNGIVEPGEECDCGP
ADAM9    375 GAS--GSRNFSSCSAEDFEKLTLNKGGNCLLNIPKPDEAYSAPSCGNKLVDAGEECDCGT
ADAM12   383 STGYPFPMVFSSCSRKDLETSLEKGMGVCLFNLPEVRESFGGQKCGNRFVEEGEECDCGE
ADAM19   375 ATGHPFPKVFNGCNRRELDRYLQSGGGMCLSNMPDTRMLYGGRRCGNGYLEDGEECDCGE
ADAM33   316 ATGHPFPRVFSACSRRQLRAFFRKGGGACLSNAPDPGLPVPPALCGNGFVEAGEECDCGP
```

```
                                                  Cysteine loop (C438-P459)
ADM-2   398 -LKCD-NHCCNGSTCKLIGEAECASGDCCDLKTCKPKPRATVCRAAIGICDLDEYCNGET
ADAM9   433 PKECELDPCCEGSTCKLKSFAECAYGDCCK--DCRFLPGGTLCRGKTSECDVPEYCNGSS
ADAM12  443 PEECM-NRCCNATTCTLKPDAVCAHGLCCE--DCQLKPAGTACRDSSNSCDLPEFCTGAS
ADAM19  435 EEECN-NPCCNASNCTLRPGAECAHGSCCH--QCKLLAPGTLCREQARQCDLPEFCTGKS
ADAM33  376 GQECR-DLCCFAHNCSLRPGAQCAHGDCCV--RCLLKPAGALCRQAMGDCDLPEFCTGTS

                       Cysteine-rich domain (C470-C611)
ADM-2   456 NDCPADFFVQNAALCPGKENEFCYEGGCGSRNDQCAKLWGPTGKNGDENCYR-KNTEGTF
ADAM9   491 QFCQPDVFIQNGYPCQNN-KAYCYNGMCQYYDAQCQVIFGSKAKAAPKDCFIEVNSKGDR
ADAM12  500 PHCPANVYLHDGHSCQDV-DGYCYNGICQTHEQQCVTLWGPGAKPAPGICFERVNSAGDP
ADAM19  492 PHCPTNFYQMDGTPCEGG-QAYCYNGMCLTYQEQCQQLWGPGARPAPDLCFEKVNVAGDT
ADAM33  433 SHCPPDVYLLDGSPCARG-SGYCWDGACPTLEQQCQQLWGPGSHPAPEACFQVVNSAGDA

ADM-2   515 HGNCGTNAHTKEIKKCETENAKCGLLQCETQAERPVFGDPGSVTFSHSTVYS-SLKRDDK
ADAM9   550 FGNCGFS-G-NEYKKCATGNALCGKLQCENVQEIPVFGI-V--P---AIIQT-PSRGTKC
ADAM12  559 YGNCGKVSK-SSFAKCEMRDAKCGKIQCQGGASRPVIGT-NAVSIE-TNIPLQQGGRILC
ADAM19  551 FGNCGKDMN-GEHRKCNMRDAKCGKIQCQSSEARPLE-S-NAVPID-TTIIM-NGRQIQC
ADAM33  492 HGNCGQDSE-GHFLPCAGRDALCGKLQCQGGKPSLLA-P-HMVPVD-STVHL-DGQEVTC

ADM-2   574 KFCYVFKSAYG----GLNAPDPGLVPDGAICGEEQMCIGQKCHKKEKISKVTA-QCLDNC
ADAM9   601 WGVD----FQL----GSDVPDPGMVNEGTKCGAGKICRNFQCVDASVL-N-YDCDVQKKC
ADAM12  616 RGTH----VYL----GDDMPDPGLVLAGTKCADGKICLNRQCQNISVF-G-VH-ECAMQC
ADAM19  606 RGTH----VYRGPEEEGDMLDPGLVMTGTKCGYNHICFEGQCRNTSFF-E-TE-GCGKKC
ADAM33  547 RGAL----ALPS--AQLDLLGLGLVEPGTQCGPRMVCQSRRCRKNAFQ-E-LQ-RCLTAC

            EGF-like (V620-V652)           Transmembrane Domain (L763-Y695)
ADM-2   629 NFRGVCNNVGNCHCERGFGGIACEIPGYGGSVNSNEAYRFRGITLSSTFLVFFCL-FGIF
ADAM9   651 HGHGVCNSNKNCHCENGWAPPNCETKGYGGSVDSGPTYNEMNTALRDGLLVFFFL-I--V
ADAM12  665 HGRGVCNNRKNCHCEAHWAPPFCDKFGFGGSTDSGPIRQADNQGLTIGILV ILC-L--L
ADAM19  659 NGHGVCNNNQNCHCLPGWAPPFCNTPGHGGSIDSGPMPPESVGPVVAGVLVAILV-L--A
ADAM33  598 HSHGVCNSNHNCHCAPGWAPPFCDKPGFGGSMDSGPVQAENHDTFLLAMLLSVLLPL--L

                Furin-2 (R696-R699)  NLS (R696-D716)  SH3 binding-1
ADM-2   688 IGG---LCVYYRVKRKRNLV-SEWWSVVKKKFDLHGDLVPVRKAPPPPYAQRIRQSFTAM
ADAM9   708 PLIVCAIFIF--IKRD-QLW----RSY---------------------FRKKRSQTYE--
ADAM12  722 AAG---FVVY--LKRK-TLI----RLL---------------------FTNKKT-TIE--
ADAM19  716 VLM---LMYY--CCRQNNKLGQLKPSA---------------------LPSKLRQQFSCP
ADAM33  656 PGA---GLAW--CCYR-LP-----------------------------------------

ADM-2   744 ------WGEDHSH--VA----VAQPAHPRNCYN--SCCRQP-PRFDPPSIPMVTLKNPNL
ADAM9   738 -----SDGKNQAN---------------------------P-SR-QPGSVP-RHVSPVTP
ADAM12  748 ----KL---RCVRPSRPPRGFQPCQAHLGHLGK--GLMRKPPDSYPPKDNPRRLLQCQNV
ADAM19  750 FRVSQNSGTGHANPTFK----LQTPQGKRKVINTPEILRKP-SQPPPR-PPPDYLRGGSP
ADAM33  669 -------------------------GAHLQRCSW--GCRRDP-ACSGPKDGPHRD------

ADM-2   789 --ASPTPLLNPAEKEEQNQERA------------THQHVELYPVAESFRSDSAASFNTRT
ADAM9   763 --PREVP----------IYANRFAVP----------TYAAKQPQQFPSRPPPPQPKVSS-Q
ADAM12  799 DISRPLNGLNVP--QPQSTQRVLPP----------LHRAPRAPSVPARPLPAKPALRQAQ
ADAM19  804 --PAPLP----------AHLSRAARNSPGPGSQIERTESSRR--PPPSRPIPPAPNCIVSQ
ADAM33  696 ---HPLGG---------------------------VHPMELGPTATGQPWPLDPENSH-E
```

```
                   SH3 binding-2                    SH3 binding-3
ADM-2    835 GSFRPNVQPPPVPRPSDDV--LSKLNEDLAKEKNAKFDRLNKTLPLPPPLPKEKPKTASS
ADAM9    801 GNL---I--PARPAPAPPLYSSLT------------------------------------
ADAM12   847 GTCKPNP--PQKPLPADPLARTTRLTHALARTPG-QWETGLRLAPL--------------
ADAM19   851 DFSRPRP--PQKALPANPVPGRRSLPRPGG------------ASPL--------------
ADAM33   725 PSS---H--PEKPLPAV-------------------------------------------

                            SH3 binding(?)
ADM-2    893 TSLRRNESIRPEQAPPPPPPAHAKPT--LPTKQPKVSEDAAATEEKVDVRSMAA------
ADAM9        ------------------------------------------------------------
ADAM12   890 ----------------R----------------PA-------------------------
ADAM19   883 ----------------RPPGAGPQQSRPLAALAPKVSPREA-LKVKAGTRGLQGGRCRVE
ADAM33   737 ----------------S----------------PD-------------------------

ADM-2    945 ---------IF-----DQKLKK---------
ADAM9        -------------------------------
ADAM12   893 -----------PQYP-HQVPRSTHTAYIK--
ADAM19   926 KTKQFMLLVVWTELP-EQKPRAKHSCFLVPA
ADAM33   740 -----------PQADQVQMPRSCLW------
```

**S2 Fig. Alignment of *C. elegans* ADM-2 with human ADAMs**

Peptide alignment of *C. elegans* ADM-2 with human meltrin family members (ADAM9/12/19/33). Predicted domains of ADM-2 are color coded. NLS, nuclear localization domain.

**S3 Fig**

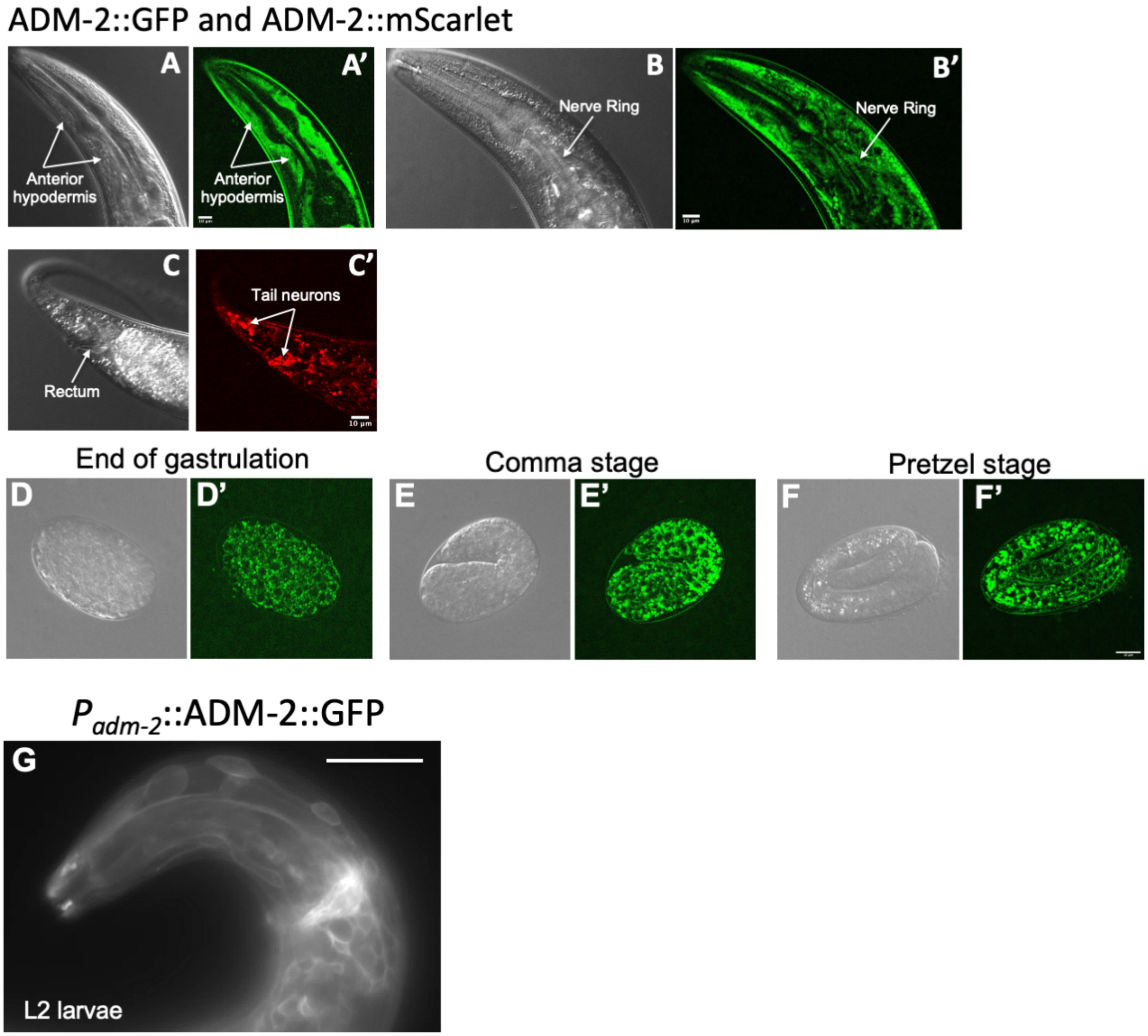


**S3 Fig. Additional ADM-2 expression**
(A–F and A’–F’) Representative DIC (A–F) and confocal (A’–F’) images of ADM-2 expression showing the anterior hypodermis (A, A’), nerve ring (B, B’), tail neurons (C, C’), and various stages of embryonic development (D–F’). Bar in A’ = 10 µm (for A, A’); in B’ = 10 µm (for B, B’); in C’ = 10 µm (for C, C’); in F’ = 10 µm (for D–F’). (G) Representative confocal image of an L2 larva expressing multi-copy $P_{adm-2}$::ADM-2::GFP in the plasma membrane of head neurons. Bar in G = 25 µm.

### S4 Fig

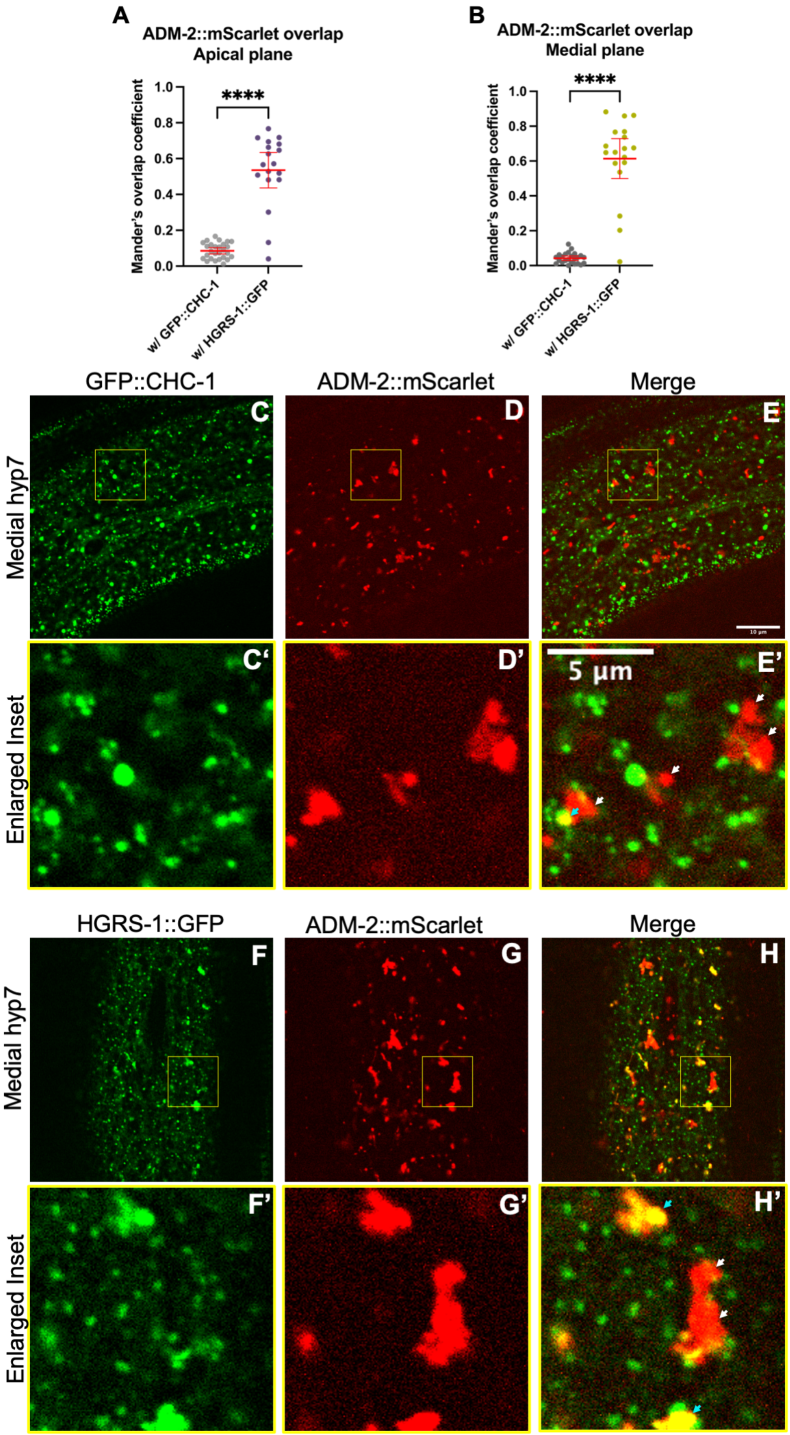


**S4 Fig. Supplemental colocalization data**

(A,B) Dot plots showing quantification of Mander's overlap coefficient for the overlap of ADM-2::mScarlet with GFP::CHC-1 and $P_{hyp7}$::HGRS-1::GFP proteins within the apical (A) and medial (B) planes. Mean values and 95% confidence intervals (error bars) are indicated. p-Values were calculated using an unpaired test: ****p ≤ 0.0001, **p ≤ 0.01. (C–H) Representative confocal images of GFP::CHC-1 (C), $P_{hyp7}$::HGRS-1::GFP (F), and ADM-2::mScarlet (D,G) within the hyp7 medial plane. C'–H' are insets of C–H confocal images. Bar in E= 10 µm (for C–H); in E' = 5 µm (for insets C'–H'). (E',H') White arrows show ADM-2 large vesicular structures that do not colocalize with GFP::CHC-1 and $P_{hyp7}$HGRS-1::GFP puncta, which are lysosomes. Cyan arrows indicate vesicles containing ADM-2 that colocalize with GFP::CHC-1 and $P_{hyp7}$::HGRS-1::GFP.

**S5 Fig**

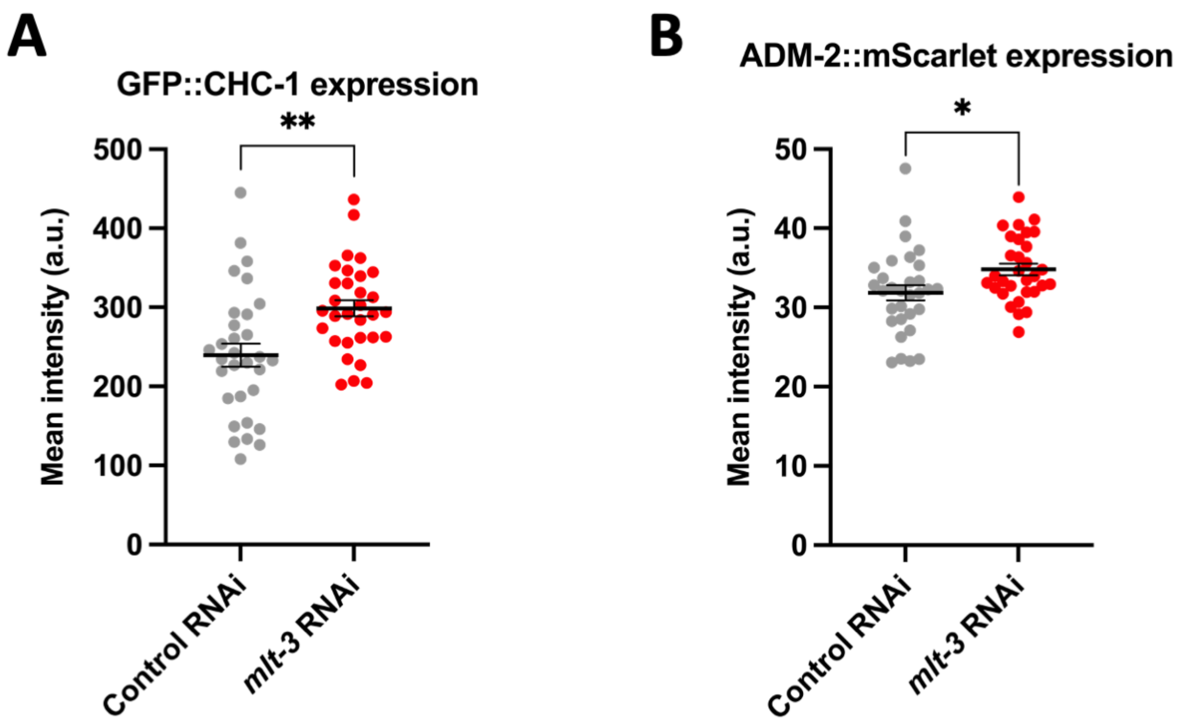


**S5 Fig. ADM-2 levels are increased slightly upon weak loss of *mlt-3***
(A,B) Dot plot showing the mean intensity (a.u.) of GFP::CHC-1 (A) and ADM-2::mScarlet (B) expression in the presence of Control RNAi (i.e., empty vector) and *mlt-3* RNAi. Group means along with 95% confidence intervals (error bars) are indicated. p-Values were obtained by comparing means using an unpaired t-test: $^{**}p \leq 0.01$, $^{*}p \leq 0.05$.

**S6 Fig**

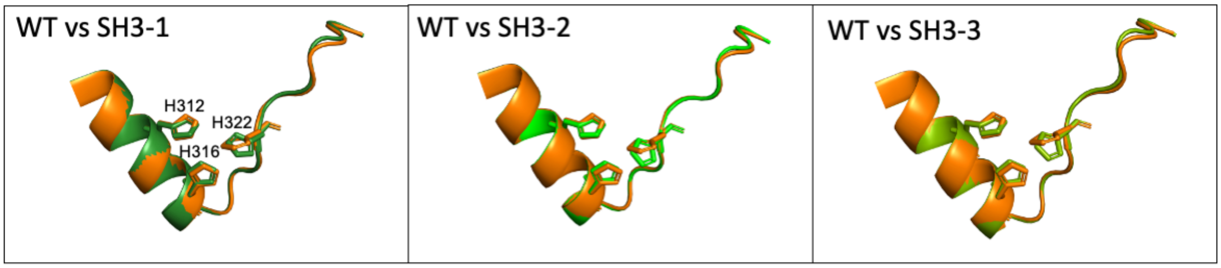


**S6 Fig. Mutations in the C-terminal region of ADM-2 do not cause conformational changes to the metalloprotease active site**
Three-dimensional protein structure of the region including amino acids 307–328 of wild-type ADM-2 (orange) superimposed on modeled structures of SH3-1, SH3-2, and SH3-3 mutant proteins.

**S7 Fig**

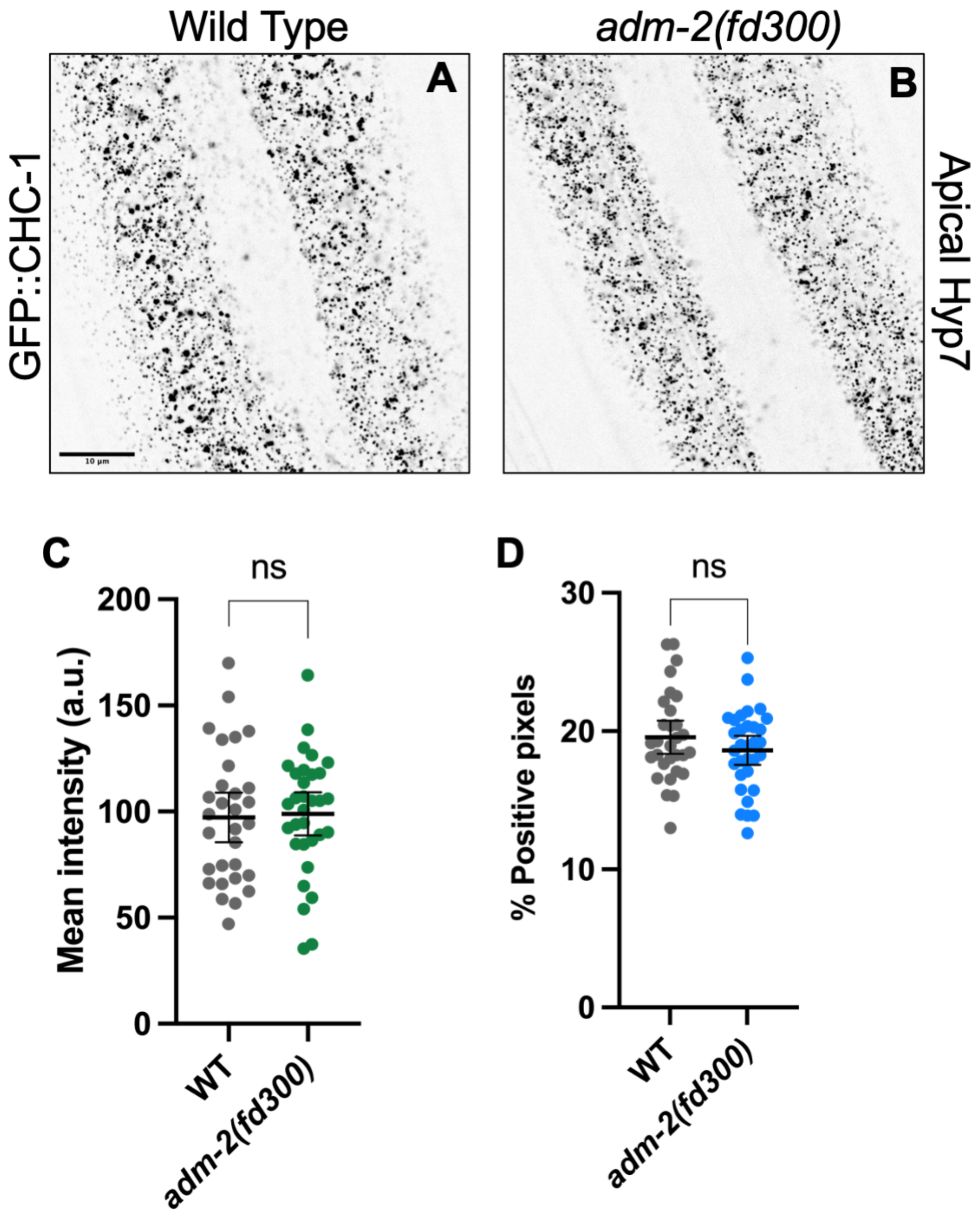


**S7 Fig. Clathrin expression is not perturbed by loss of ADM-2 function**
(A,B) Representative confocal images of GFP::CHC-1 expression in the hyp7 region of the hypodermis in the apical plane in wild-type (A) and *adm-2(fd300)* null mutant (B) 1-day-old adult worms. Bar in A = 10 µm (for A,B). (C,D) Dot plots showing GFP::CHC-1 mean intensity (a.u.) (C) and the percentage of GFP-positive pixels (D) within the apical plane for each individual worm of the specified genotype. In C and D, group means along with 95% confidence intervals (error bars) are indicated. p-Values were obtained by comparing means using an unpaired t-test. ns, $p > 0.05$.
