## Supplementary material for "An unexpected role for the conserved ADAM-family metalloprotease ADM-2 in *Caenorhabditis elegans* molting": S1 Table. List of strains

**Table S1. Strains used in this study**

| Strain | Genotype |
|---|---|
| RT3402 | *pw17(gfp::chc-1)* |
| WY1145 | *nekl-2(fd81); nekl-3(gk894345); fdEx286 (pDF153(nekl-3 (+)); pTG96(sur-5::GFP))* |
| WY1208 | *nekl-2(fd81); nekl-3(gk894345) adm-2(fd130)* |
| WY1232 | *nekl-2*(*fd81*); *nekl-3*(*gk894345*); *fdEx186* (*nekl-3*$^{+}$ + SUR-5::GFP); *fdEx197* (SUR-5::RFP)] |
| WY1279 | *nekl-2(fd81); nekl-3(gk894345) adm-2(fd163)* |
| WY1347 | *nekl-2(fd81); nekl-3(gk894345) adm-2(fd208)* |
| WY1342 | *nekl-2(fd81); pwIs528(GFP::CHC-1); nekl-3(gk894345) adm-2(fd130); fdEx286* |
| WY1386 | *nekl-2(fd81); pwIs528(GFP::CHC-1); nekl-3(gk894345) adm-2(fd130); fdEx286; fdEx315 (adm-2 fosmid mix; WRM0620dD12, WRM0632aG02, and WRM0610cA04)* [Line #1] |
| WY1388 | *nekl-2(fd81); pwIs528(GFP::CHC-1); nekl-3(gk894345) adm-2(fd130); fdEx286; fdEx356 (adm-2 fosmid mix; WRM0620dD12, WRM0632aG02, and WRM0610cA04)* [Line #2] |
| WY1428 | *nekl-2(fd81); nekl-3(gk894345) adm-2(fd228)* [large deletion] |
| WY1429 | *nekl-2(fd81); nekl-3(gk894345) adm-2(fd229)* [large deletion] |
| WY1430 | *nekl-2(fd81); nekl-3(gk894345) adm-2(fd230)* [large deletion] |
| WY1435 | *adm-2(fd235)* [large deletion] |
| WY1436 | *adm-2(fd236)* [large deletion] |
| WY1437 | *adm-2(fd237)* [large deletion] |
| WY1605 | *pw17(gfp::chc-1); adm-2(fd300)* |
| WY1640 | *nekl-2(fd91); adm-2(fd313); fdEx278* [pDF166 (*nekl-2* genomic) + pTG96 (SUR-5::GFP)] |
| WY1643 | *nekl-3(sv3) adm-2(fd316); mnEx174* [F19H6 (*nekl-3* genomic) + pTG96 (SUR-5::GFP)] |
| WY1644 | *mlt-4(sv9); adm-2(fd317) mlt-4(sv9); mnEx173* [ZC15 (*mlt-4* genomic) + pTG96 (SUR-5::GFP)] |
| WY1562 | *eqIs1(lrp-1::gfp); ieSi57(peft-3::mRuby::tir-1); pw29(nekl-3::aid)* |
| WY1654 | *eqIs1(lrp-1::gfp); fcho-1(ox477::unc-119(+)); ieSi57(peft-3::mRuby::tir-1); pw29(nekl-3::aid)* |
| WY1655 | *eqIs1(lrp-1::gfp); adm-2(fd318); ieSi57(peft-3::mRuby::tir-1); pw29(nekl-3::aid)* |
| WY1656 | *eqIs1(lrp-1::gfp); adm-2(fd300)* |
| LH191 | *eqIs1(lrp-1::gfp); rrf-3(pk1426)* |
| WY1657 | *adm-2(fd300)* |
| PHX2391 | *adm-2::eGFP (syb2391)* |
| PHX1722 | *adm-2::mScarlet (syb1722)* |
| WY1664 | *pw17(gfp::chc-1); adm-2::mScarlet (syb1722)* |
| WY1675 | *adm-2::mScarlet (syb1722); pwSi125[phyp7::NeonGreen::hgrs-1]* |
| WY1820 | *N2; fdEx373* |
| WY1821 | *N2; fdEx374* |
| WY1822 | *N2; fdEx375* |
| WY1833 | *N2; fdEx382* |
| WY1841 | *eqIs1(lrp-1::gfp); fdEx382* |
| WY1893 | *pw27(nekl-2::aid); ieSi10(phyp7::BFP::tir-1); adm-2::mScarlet (syb1722)* |
| WY1897 | *ieSi10(phyp7::BFP::tir-1); pw29(nekl-3::aid) adm-2::mScarlet (syb1722)* |
| WY1431 | *nekl-2(fd81); nekl-3(gk894345) adm-2(fd231)* |

| | |
|---|---|
| WY1513 | *nekl-2(fd81)*; *nekl-3(gk894345) adm-2(fd243)* |
| WY1585 | *nekl-2(fd81)*; *nekl-3(gk894345) adm-2(fd288)* |
| WY1518 | *nekl-2(fd81)*; *nekl-3(gk894345) adm-2(fd248)* |
| WY1522 | *nekl-2(fd81)*; *nekl-3(gk894345) adm-2(fd252)* |
| WY1527 | *nekl-2(fd81)*; *nekl-3(gk894345) adm-2(fd257)* |
| WY1676 | *nekl-2(fd81)*; *nekl-3(gk894345) adm-2(fd324)* |
| WY1669 | *nekl-2(fd81)*; *nekl-3(gk894345) adm-2(fd322)* |
